# Characterizing the Microbiome of ’Sterile’ Organs in Experimental Mice

**DOI:** 10.1101/2025.04.02.646907

**Authors:** Ming Xu, Shuyun Guan, Chaoran Zhong, Mingyang Ma, Li Tao, Guanghua Huang

**Affiliations:** State Key Laboratory of Genetic Engineering, School of Life Sciences, Department of infectious diseases, Department of Laboratory Medicine, Huashan Hospital, Shanghai Institute of Infectious Disease and Biosecurity, Fudan University, Shanghai 200438, China; College of Pharmaceutical Sciences, Southwest University, Chongqing 400716, China

**Keywords:** Experimental mice, microbiota, culturomics, metagenomics, sterile organs

## Abstract

Recent studies have shown that microorganisms can be present in organs or blood traditionally considered sterile, both in healthy and diseased individuals. In this study, we investigated the presence of microorganisms in the “sterile” organs (brain, heart, kidney, liver, lung, and spleen) of commercially available experimental mice using culturomics and metagenomics. To avoid potential environmental contamination, strict disinfection assays were performed. Among 104 mice of the C57BL/6J, BALB/c, and ICR strains purchased from three major experimental animal suppliers in China, 23 mice (22.1%) exhibited a high microbial burden (>1 x 10^4^ CFU/g tissue) in at least one organ, and 42 mice (40.4%) exhibited >1 x 10^3^ CFU/g tissue in at least one organ. In total, 216 microbial species were identified through culturomics. Of them, 56 microbial species were found in the organs of at least three mice of the 42 mice (>1 x 10^3^ CFU/g tissue in at least one organ). Metagenomics analysis of 10 high-burden mice verified the existence of microbes in the mouse organs and identified 262 species. Several species, including *Acinetobacter* sp., *Alcaligenes faecalis*, *and Ligilactobacillus murinus*, were identified as the most abundant microbes across both culturomics and metagenomics assays. 16S FISH staining assays verify the presence of microbes in the mouse organs. Feces Metagenomics analysis and detection of the wide distribution of *L. murinus* in the mouse mesenteric lymph nodes (MLNs) imply that microbial cells could be translocated from the gut to the MLNs and then to the remote tissues. Together, these findings reveal the presence of non-pathogenic microorganisms in the “sterile” tissues of experimental mice, raising concerns about potential variability in experimental outcomes when using such animals in research.

## Introduction

Myriads of microorganisms, collectively referred to as the microbiome or microbiota, colonize the skin, mucosal surfaces, and the digestive and genital tracts of mammals, including humans [1–5]. The microbiome has significant positive and negative effects on human health, influencing processes such as nutrient absorption, immune system development and regulation, infection prevention, and diseases progress [6–14]. Traditionally, internal organs and tissues, including the brain, heart, liver, spleen, and kidney, were believed to be sterile [15]. However, recent studies have challenged this notion, demonstrating that the presence of living microorganisms in the blood and other organs of both healthy and diseased individuals [16–21]. While the physiological roles of these intratissue microbes remain unclear, their discovery reshapes our understanding of host-microbe interactions.

Evidence suggests that intratumor microbiomes may influence the initiation, progression, metastasis, and therapeutic responses of various cancers [22–33]. For instance, unique microbial signatures have been identified in the blood and tissues of cancer patients [20] [28]. Leinwand *et al*. (2022) recently reported the presence of microorganisms in the liver and their role in modulating hepatic immunity and tolerance [34]. Intratissue or intratumor microbes may originate from adjacent normal tissues (NATs) [35] or result from gut bacterial translocation, as observed in pancreatic ductal adenocarcinoma [17, 36, 37].

Microbiomes also play critical roles in the health of animals [38]. Experimental animals, such as mice and rats, are widely used in biological and medical research [39]. To ensure reproducibility, inbred strains of laboratory animals are commonly employed, and their feeding and housing conditions are standardized in specific-pathogen free (SPF) research facilities. Despite these controlled conditions, significant variations in experimental outcomes are often observed among individual animals [40]. These variations are thought to be influenced by uncontrollable environmental factors, particularly microbiota [41, 42]. For example, differences in fungal burdens and survival times have been reported among mice in systemic infection models under tightly regulated conditions [43, 44]. Consistent with these investigations, researchers have detected living microorganisms in the liver and low levels of bacterial 16S rDNA sequences in the “sterile” organs of laboratory mice or rats [45, 46]. Laboratory mice, such as C57BL/6J, BALB/c, and ICR strains, are not true “wild-type” animals. Long-term inbreeding has led to the accumulation of genomic or phenotypic deficiency [47–49]. For example, C57BL/6J mice carry natural mutations in exons 7-11 of *Nnt* (a gene encoding the nicotinamide nucleotide transhydrogenase), while BALB/c mice are known for their extreme susceptibility to infections and carcinogens [50, 51]. These genetic deficiencies may compromise immune system function, potentially facilitating microbial translocation from the gut to other tissues [52]. A similar phenomenon could occur in experiment mice due to their genetic predispositions.

Given the extensive use of laboratory mice in biomedical research, it is crucial to determine whether living microorganisms are present in organs traditionally considered “sterile” and assess their potential effects on animal physiology and experimental outcomes. In this study, we used culturomics and metagenomics approaches to investigate the microbiomes of “sterile” organs in 104 mice from C57BL/6J, BALB/c, and ICR strains, sourced from multiple experimental animal providers. We found that over 20% of the mice harbored significant microbe burdens (>1 x 10^4^ CFU/g tissue) in the brain, heart, kidney, liver, lung, and spleen tissues. While microbial diversity varied among individual animals, we identified a core microbiome. Metagenomics analysis further revealed a potential link between the microbiomes of “sterile” organs and feces. These findings highlight the variability in microbiomes among laboratory mice and underscore the importance of considering these factors when interpreting experimental outcomes.

## Results

### Concept and research design for microbiome analysis of experimental mouse “sterile” organs

Emerging evidence suggests the presence of microorganisms in traditionally considered “sterile” organs in both healthy individuals and those with diseases such as cancers [34–37]. These microorganisms may exert positive or negative effects on health, physiology, disease progression, or therapeutic outcomes. This study aimed to determine whether microorganisms exist in the “sterile” organs of experimental mice, a widely used model in biological and biomedical research. We also investigated whether these microorganisms could affect research outcomes and sought to draw parallels with microbiomes in other healthy animals or humans. To address these questions, we designed culturomics and metagenomics experiments to identify viable microorganisms in the brain, heart, kidney, liver, lung, and spleen tissues of mice (**Figure 1A**).

**Figure 1.**
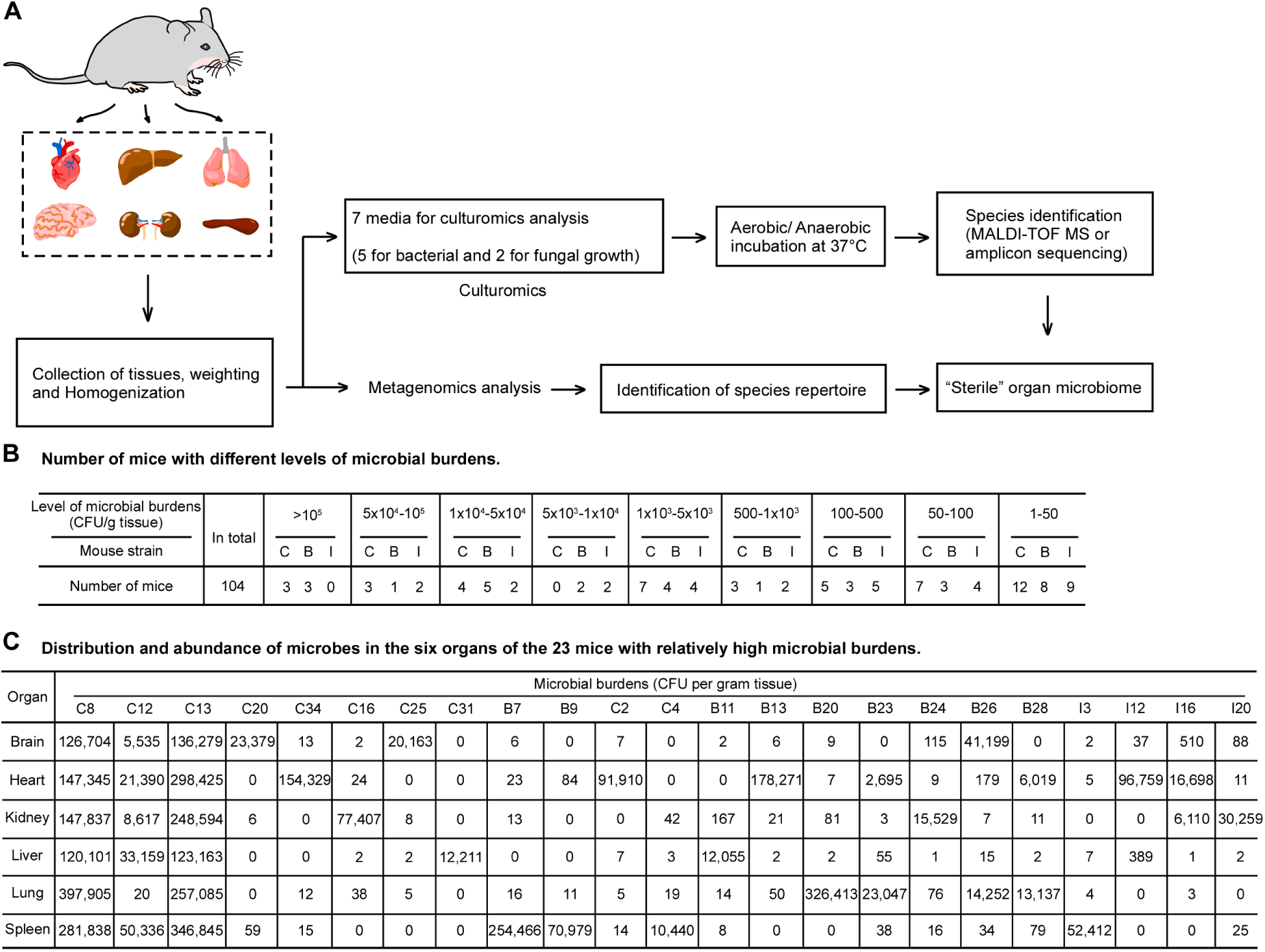
Summary of the workflow and outcomes of this study. (A) A schematic workflow. A total of 104 mice from three strains (C57BL/6J, BALB/c, and ICR), purchased from three major experimental animal providers (WTL, NMO, and ZYU), were subjected to culturomics and metagenomics analyses (**Table S1**). The mice were housed in an SPF animal facility. After humane euthanasia, organs (brain, heart, kidney, liver, lung, and spleen) were collected immediately for culturomics and/or metagenomics analyses. For culturomics, the seven most optimal cultural media, previously tested, were selected for bacterial and fungal isolation growth under aerobic or anaerobic conditions. Two representative colonies of each distinct colony morphology on each plate were selected for species identification. For metagenomics, organs from ten mice with a relatively high microbial burden were analyzed using 2bRAD-M sequencing. (B) Number of mice with different levels of microbial burdens in the “sterile” organs. The figure shows the microbial burden levels in at least one of six tissues (brain, heart, kidney, liver, lung, and spleen). CFU, colony-forming units. CFU levels (CFU/g tissue), mouse strains, and the number of mice with varying microbial abundance are indicated. (C) Distribution and abundance of microbes in the six organs of 23 mice with relatively high microbial burdens (> 10^4^ CFU/g tissue) detected by culturomics. The figure shows the microbial abundance (CFU/g tissue) in the six organs of 23 high burden mice. Mouse strains: C, C57BL/6J; B, BALB/c; I, ICR. This figure is associated with Figures 3**, S1 and Datasets S1** and **S2**.

A total of 104 mice (44 C57BL/6J, 30 BALB/c, and 30 ICR) from three Chinese providers (WTL, NMO, and ZYU) were included (**Figure 1B** and **Table S1**). All mice were healthy with no observed debilitating phenotypes. Mice were humanely euthanized, strictly disinfected with 75% ethanol and dissected under sterilized conditions. The surface of the dissected tissues was washed carefully with 75% ethanol and then rinsed with sterile PBS three times. In total, 624 organ samples were collected for microbiome analyses, which included both culturomics and metagenomics approaches. Parallel environmental negative controls were performed to rule out contamination originating from the laboratory. After screening 30 culture media for culturomics, we selected 7 optimal media (BHI1, BHI2, AIA, Chocolate agar, YCFA, YPD and CDA) for bacterial and fungal growth under aerobic and anaerobic conditions at 37°C (**Table S2**). YPD and CDA media were used for fungal isolation. Culturomics analysis was performed on all 104 mice. The dominant microbial species in different organs were analyzed in 23 mice with ≥1 x 10^4^ CFU/g tissue in at least one organ (**Figure 1C**), while metagenomics analysis was conducted on 10 mice with the highest microbial abundance identified by culturomics (mouse code: C8, C12, C13, C16, C20, C25, C31, C34, B7, and B9).

### Culturomics assays identify viable microorganisms in the mouse organs

As shown in **Figure 2A**, distinct microbial colonies formed on the plates when ground tissues were plated onto media and incubated under either aerobic or anaerobic conditions. To confirm the presence of microbes in the animal organs, we performed FISH assays based on 16S rRNA hybridization, which revealed clear fluorescence signals in all tested tissues (**Figure 2B**). The microbial burden (CFU) in the brain, heart, kidney, liver, lung, and spleen tissues of each mouse was assessed by quantifying colonies on seven optimal media (**Figures 1B**, **3**, and **S1** and **Dataset S1**). Among the 104 mice tested, 6 (5.77%) exhibited microbial loads exceeding 1 x 10^5^ CFU/g tissue, 17 (16.35%) had loads between 1 x 10^4^ to 1 x 10^5^ CFU/g, and 19 (18.27%) had loads ranging from 1 x 10^3^ to 1 x 10^4^ CFU/g in at least one organ (**Figures 1B**, **3 and S1**). Notably, 8 mice exhibited CFU values exceeding 1 x 10^3^ in at least two organs (**Figures 3 and S1**). For the C57BL/6J mice, 18 mice had relatively low microbial burden, with CFU values below 1 x 10^2^ CFU/g across all six organs, while 8 had CFU values ranging from 1 x 10^2^ to 1 x 10^3^ CFU/g in at least one organ. In contrast, 11 mice (45.8%) from the WTL provider, 5 (50%) from the NMO provider, and 1 (10%) from ZYU provider exhibited CFU values exceeding 1 x 10^3^ CFU/g in at least one organ. Two mice (C8 and C13) showed high CFU values (>1 x 10^5^ CFU/g) across six organs, while one mouse (C12) had similar high CFU values (>1 x 10^5^ CFU/g) in three organs. Additionally, two mice (C2 and C34) exhibited high CFU values (∼ 1 x 10^5^ CFU/g) in the heart, and two others (C20 and C25) had elevated CFUs (>1 x 10^4^ CFU/g) in the brain. Despite the high microbial loads in some organs, CFU values in mouse blood were consistently very low (**Dataset S1**). Furthermore, to determine whether the organ microbiome changes with sex or age, we comparatively analyzed the organ microbial burden in both male and female mice aged 4, 6, 7 weeks, or 3, 12, 21 months, respectively (**Table S1**). However, no clear association was observed between organ microbial abundance and the sex or age of the mice (**Figure 3 and Dataset S1**). Further studies are needed to address these questions.

**Figure 2.**
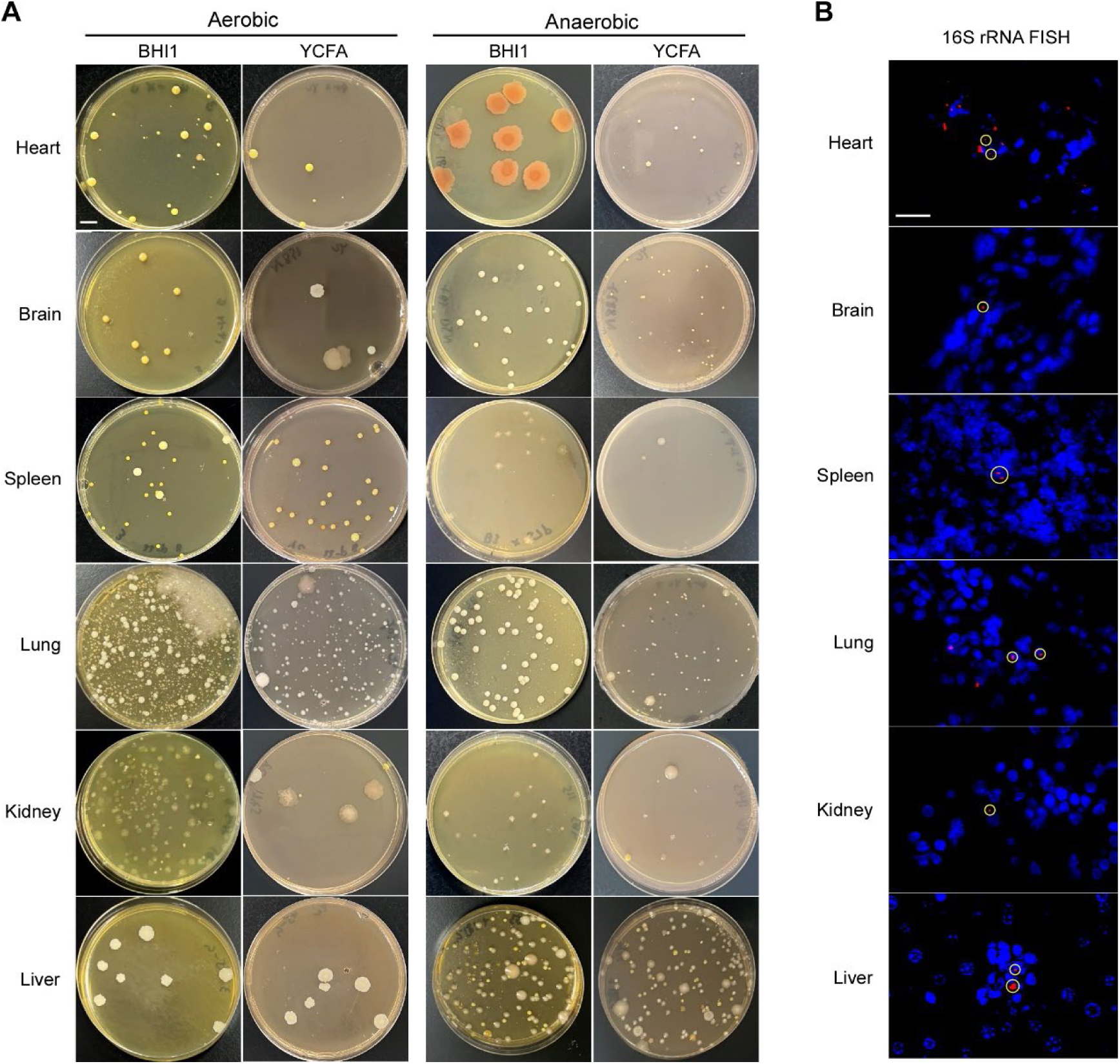
Representative examples of culture plates and FISH images for microbial detection in mouse brain, heart, kidney, liver, lung, and spleen tissues. (A) Microbial colonies on BHI1 and YCFA medium plates. After the mice were humanely euthanized, their organs (brain, heart, kidney, liver, lung, and spleen) were immediately used for culturomics analysis. The plates were cultured under aerobic or anaerobic condition at 37 °C for 5 days. Scale bar: 1 cm. (B) Representative images for fluorescence in situ hybridization (FISH) assays using the bacterial 16S rRNA probe in mouse tissues. The dissected tissues were fixed with 10% formalin, embedded in paraffin, and sectioned. Tissue sections were hybridized with Cy3-labeled probe EUB338 (5’-GCTGCCTCCCGTAGGAGT-3’) (red) and 6-FAM labeled nonspecific complement probe (5’-CGACGGAGGGCATCCTCA-3’) (green). The latter was used as a negative control to rule out non-specific hybridization. After hybridization, the sections were washed to remove the hybridization solution, and nuclei were counterstained with diamidino-phenyl-indole (DAPI). The images were captured using fluorescent microscopy. FAM (488) glows green by excitation wavelength 465-495 nm and emission wavelength 515-555 nm; CY3 glows red by excitation wavelength 510-560 nm and emission wavelength 590 nm. Circles indicate bacteria stained by the FISH probe. Scale bar: 20 μm. Representative images for each tissue are shown.

**Figure 3.**
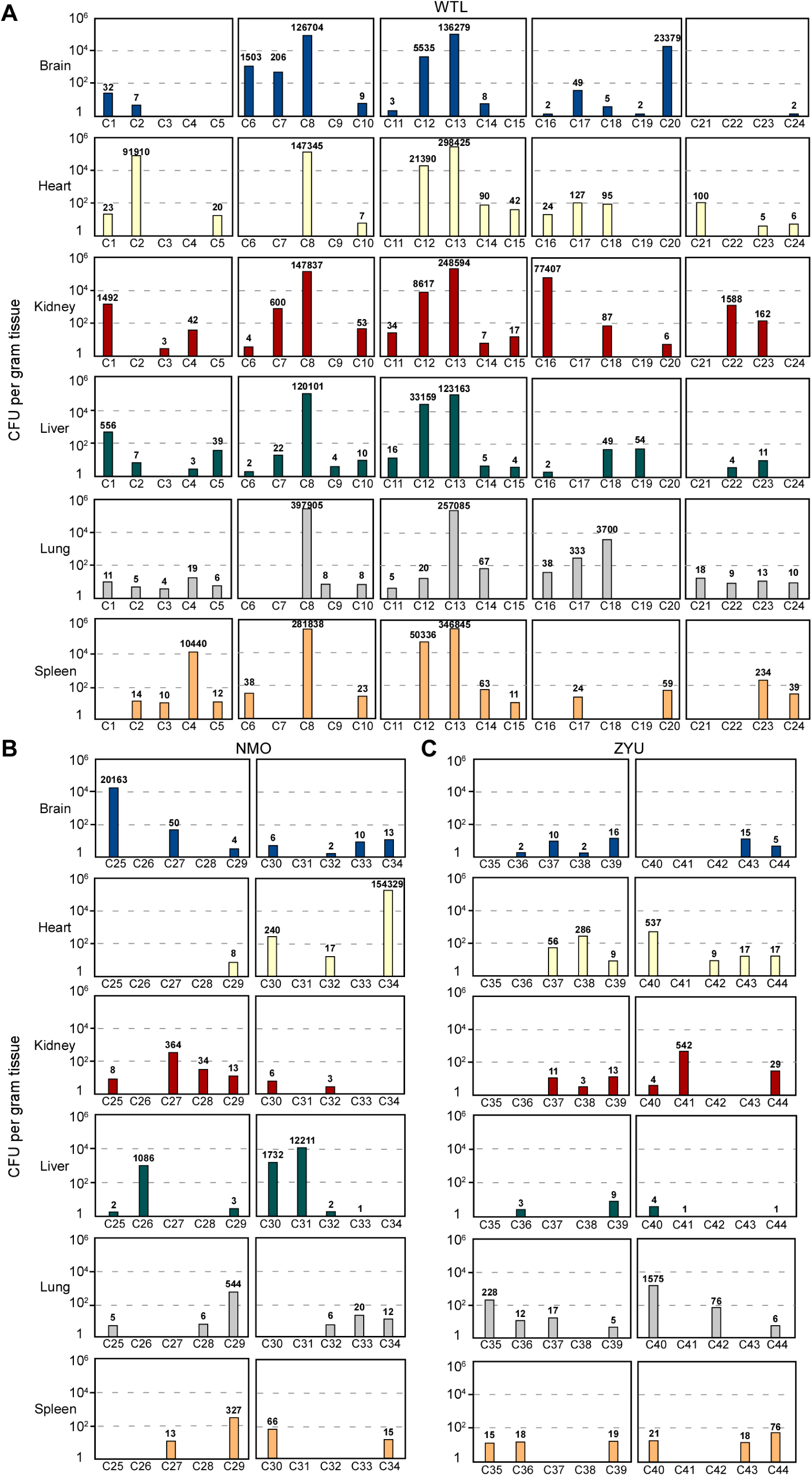
Abundance of microbes detected in the brain, heart, kidney, liver, lung, and spleen tissues of C57BL/6J mice based on culturomics assays. Microbial burdens were evaluated in a total of 44 C57BL/6J mice purchased from three experimental animal providers in China: WTL (A), NMO (B), and ZYU (C). Upon arrival, the mice were immediately euthanized, and their organs were used for microbial burden analysis. After complete anesthesia, the body surface was sterilized three times with 75% ethanol. The microbial abundance (CFU/g tissue) in the six organs was evaluated at 37^°^C using seven optimized cultural media. CFU, colony-forming units. The column colors represent specific organs, with the numbers on the columns indicating CFU values. Detailed information about the mice used is provided in **Table S1**. C1-C44, mouse codes ranked by treatment order. C: C57BL/6J.

Similar microbial burdens were observed in the organs of two other mouse strains, BALB/c and ICR, obtained from the WTL, NMO, and ZYU commercial experimental animal providers (**Figure S1 and Dataset S1**). Notably, the microbial loads in C57 and ICR mice purchased from ZYU were lower compared to those from WTL and NMO. Collectively, these findings demonstrate the presence of viable microbes in the organs traditionally considered “sterile” in experimental mice. However, microbial abundances varied across organs and individual mice depending on the commercial providers, mouse strains, or batches.

### Distribution and abundance of microbial species in mouse organs revealed by culturomics analysis

From the culture plates representing culturomics results of the “sterile” organs of 104 mice, a total of 2,600 colonies were selected for species identification. Two representative colonies from each colony phenotype grown on individual medium plate were analyzed using MALDI-TOF assays. For colonies that could not be accurately identified due to technical limitation, molecular identification through amplicon sequencing was performed. In total, 216 microbial species were identified, including 213 bacterial and 3 fungal species (**Dataset S1**). The composition of microbial species varied significantly among individual mice. Three mice (C3, C31, B6) harbored only a single microbial species across their six organs, while 25 mice carried between 2 to 5 species, and the remaining mice hosted more than 5 species. Notably, mice B11, B26, and I12 exhibited the highest diversity, harboring 28, 24, and 21 microbial species, respectively. Of the total 216 microbial species, 56 (25.9%) were detected in at least three mice of the 42 mice with a relatively high microbial burden (>1 x 10^3^ in at least one organ) (**Figure 4**). For example, *Staphylococcus epidermidis* was detected in 23 mice (54.8%), *Micrococcus luteus* in 20 mice (47.6%), *L. murinus* in 16 mice (38.1%), *Staphylococcus hominis* and *Bacillus cereus* in 15 mice (35.7%), *Micrococcus endophyticus* in 11 mice (26.2%), *Microbacterium* sp. in 10 mice (23.8%) (**Figure 4, Dataset S2**). Among the 56 bacterial species, 45 (80.4%) were Gram-positive, including 26 *Bacillota*, 16 *Actinomycetota*, and 3 *Actinobacteria*. The remaining 11 (19.6%) Gram-negative species belong to the phylum *Pseudomonadota* (**Dataset S2**). Phylogenetic analysis based on 16S rDNA sequences revealed that high-abundant species were distributed across three phyla: *Pseudomonadota, Actinomycetota*, and *Bacillota* (**Figure S2 and Dataset S2)**. For example, the *Pseudomonadota* phylum included *Alcaligenes faecalis*, *Brevundimonas diminuta*, and *Acinetobacter* sp., while the *Actinomycetota* phylum contained *Microbacterium* sp. and *Micrococcus* sp. The Bacillota phylum included high-abundance species such as *Staphylococcus* sp., *Bacillus* sp., and *L. murinus*.

**Figure 4.**
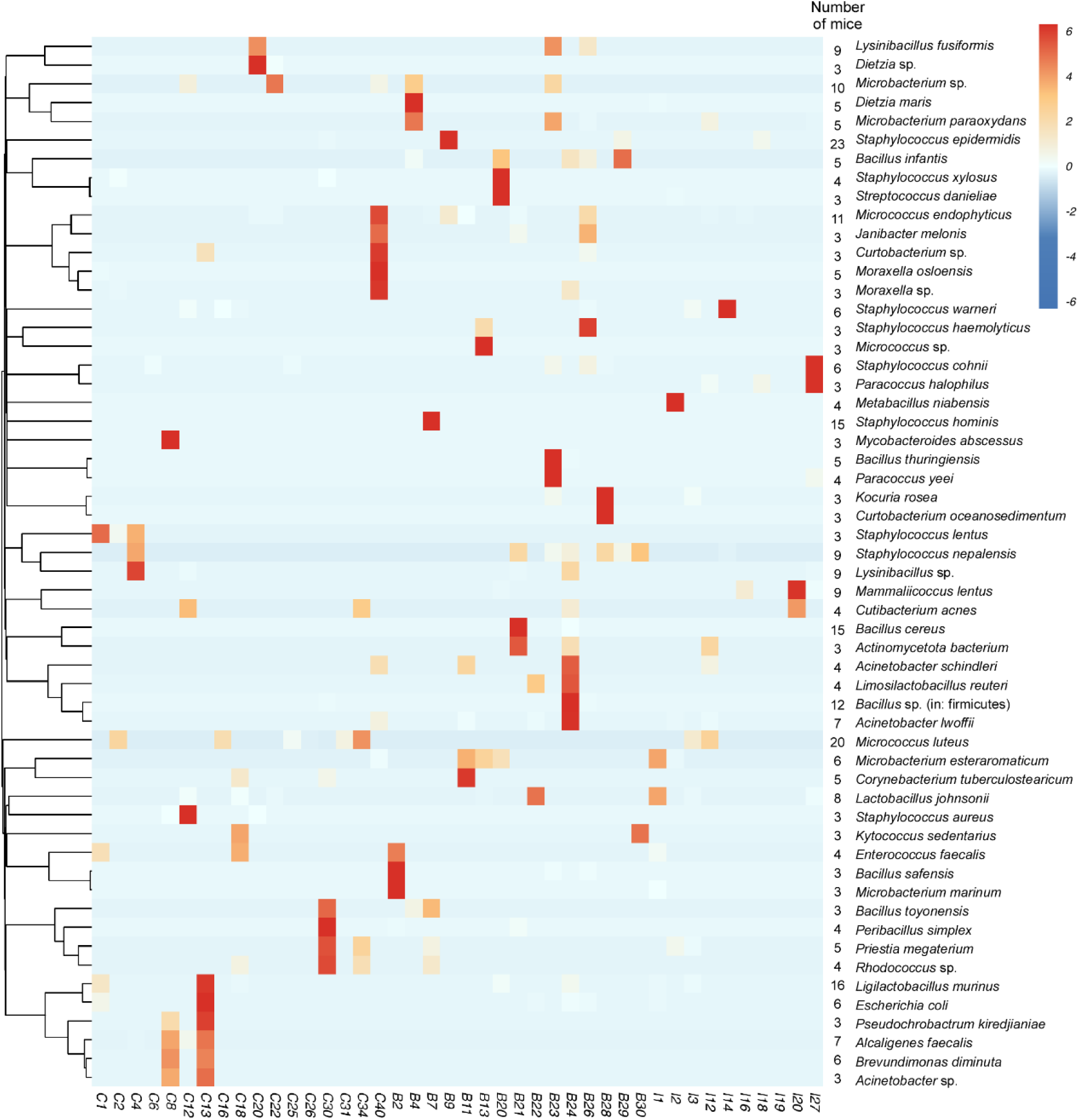
Composition and abundance of microbial species isolated by culturomics assays from six organ tissues of mice with a high microbial burden. 42 mice with a microbial burden higher than 1 x 10^3^ CFU/g tissue in at least one organ were detected. The heatmap depicts the composition and abundance of microbial species detected in the organs of at least three mice. Microbial species and number of mice harboring the corresponding species were displayed on right side of the heatmap. For species identification, two representative colonies of each distinct colony morphology on each plate were selected. The CFU of each microbial species in an organ was estimated based on similar colony phenotype on the same plate. The heatmap, representing relative abundance, was calculated as the average CFU/g tissue across the six organs. The heatmap representing relative abundance, was generated using OECloud tools (https://cloud.oebiotech.com). The intensity of red coloration indicates the relative microbial abundance. The dendrogram illustrates the phylogenetic relationships of the microbial cohorts. Mouse codes shown at the bottom of the image correspond to the treatment order. C, C57BL/6J; B, BALB/c; and I, ICR. This figure is associated with Figures 3**, S1** and **Datasets S1, S2** and **S3**.

The distribution of dominant microbial species also varied among different mouse strains. The most frequently isolated microbial species from the organs of C57BL/6J, BALB/c, ICR mice are shown in **Figure 4, Dataset S1** and **Dataset S2**. For example, *Alcaligenes faecalis*, *Brevundimonas diminuta*, and *Pseudochrobactrum kiredjianiae* were dominant in C57BL/6J mice, while *Lysinibacillus fusiformis* and *Corynebacterium tuberculostearicum* were found in both C57BL/6J and BALB/c mice. In contrast, *Micrococcus luteus* was dominant in C57BL/6J and ICR mice, and *Staphylococcus epidermidis* was common in BALB/c and ICR mice (**Figure 4**).

Further analysis was performed to determine microbial distribution and abundance in the six organs of different mice (**Dataset S3**). An UpSet plot analysis was performed to identify species that were unique or shared among different organs based on the analysis of the 42 mice with a relatively high microbial burden (>1 x 10^3^ in at least one organ). Microbial species isolated from the same organ of at least three individual mice were analyzed. As shown in **Figure S3** and **Dataset S3**, 9 microbial species were found to be unique in a particular organ, whereas 13 species were shared across different organs. Three species, *Paracoccus yeei*, *Acinetobacter lwoffii*, and *Moraxella osloensis*, were only found in the liver, whereas *Dietzia maris*, *Limosilactobacillus reuteri*, and *Pseudochrobactrum kiredjianiae* were only found in the kidney. *Acinetobacter* sp., and *Escherichia coli* were unique in the lung, and *Curtobacterium* sp. was unique in the heart. *L. murinus*, *Staphylococcus hominis*, *Bacillus cereus*, and *Micrococcus luteus* were identified in four organs, while *Mammaliicoccus lentus*, *Lysinibacillus* sp., and *Lactobacillus johnsonii* were found in three organs. Furthermore, we analyzed the dominant species among different organs in the 23 mice with a high microbial burden (>10^4^ CFU/g tissues in at least one organ). As shown in **Figure S4,** *Alcaligenes faecalis* and *Brevundimonas diminuta* were dominant in all organs except the lung, while *Staphylococcus* sp. was dominant in all organs except the heart. *Alcaligenes faecalis* was the most dominant species in the brain, spleen, kidney, and liver. Additionally, *Micrococcus luteus* were dominant in the brain, heart, and spleen, while *Bacillus* sp. was enriched in kidney, liver, and lung (**Figure S4, Dataset S3**). Taken together, the six organs exhibited unique microbial signatures, and individual mice displayed similar distribution pattern. The dominant species identified in the 23 high-burden mice were also prevalent in a broader population. For example, *Micrococcus luteus* was found in 47.6% of the mice (20/42), *L. murinus* in 38.1% (16/42), *Mammaliicoccus lentus* in 21.4% (9/42), and *Alcaligenes faecalis* in 16.7% (7/42).

### Metagenomics analysis of the organ microbiomes of experimental mice

Since many microbial species cannot be cultured under *in vitro* conditions and culture-based methods have limitations in evaluating microbial abundance, we performed metagenomics analysis using a 2bRAD-M sequencing strategy to evaluate the microbiomes of mouse organs. We focused on the brain, heart, kidney, liver, lung, and spleen tissues from 10 mice with relatively high microbial abundance as revealed by culturomics assays. A total of 262 microbial species were identified across these six organs (**Dataset S4**). Of the 262 species identified by metagenomics, 15 species were common to both the culturomics and metagenomics assays (**Figure 5A**). Among the common species, *Acinetobacter*sp., *Alcaligenes faecalis*, *Escherichia coli*, *L. murinus*, *Microbacterium* sp., *Micrococcus luteus*, and *Pseudochrobactrum asaccharolyticum*, exhibited significantly high relative abundances in both analyses (**Figure 5B**). Metagenomics analysis distinct microbial diversity across the 10 mice. For example, mice C34 and C20 harbored 123 and 76 microbial species, respectively, while mice C25 and C31 had only 11 and 17 microbial species, respectively (**Dataset S4**). In total, 97 microbial species (37.02%) were detected with a relative abundance >10^-4^ in at least two mice. Those species were distributed across 7 phyla, 11 classes, 25 orders, 39 families, 74 genera, and 97 species. The identified phyla included *Bacillota* (30/97, 30.93%), *Pseudomonadota* (21/97, 21.65%), *Bacteroidota* (24/97, 24.74%), *Actinomycetota* (19/97, 19.59%), *Ascomycota* (1/97, 1.03%), *Deinococcota* (1/97, 1.03%), and *Desulfobacterota* (1/97, 1.03%). Despite the distinct microbial diversity observed in each mouse, several species were common across the three mice (C8, C12, and C13). For example, *Alistipes* sp., *Anaerotruncus* sp., *Duncaniella dubosii*, *Duncaniella muris*, *Eubacterium* sp., *L. murinus*, *Muribaculum arabinoxylanisolvens*, *Muribaculum intestinale*, *Paramuribaculum intestinale*, *Rothia mucilaginosa* were detected. Of these, *Alistipes* sp. and *Duncaniella dubosii* were detected in more than 8 mice (**Figure 6, Dataset S4**). Metagenomics assays identified a much broader range of microbial species compared to culturomics assays likely due to the fact that some species are difficult to culture. Furthermore, metagenomics can detect both living and dead microbes, whereas culturomics only identifies living, culturable species.

**Figure 5.**
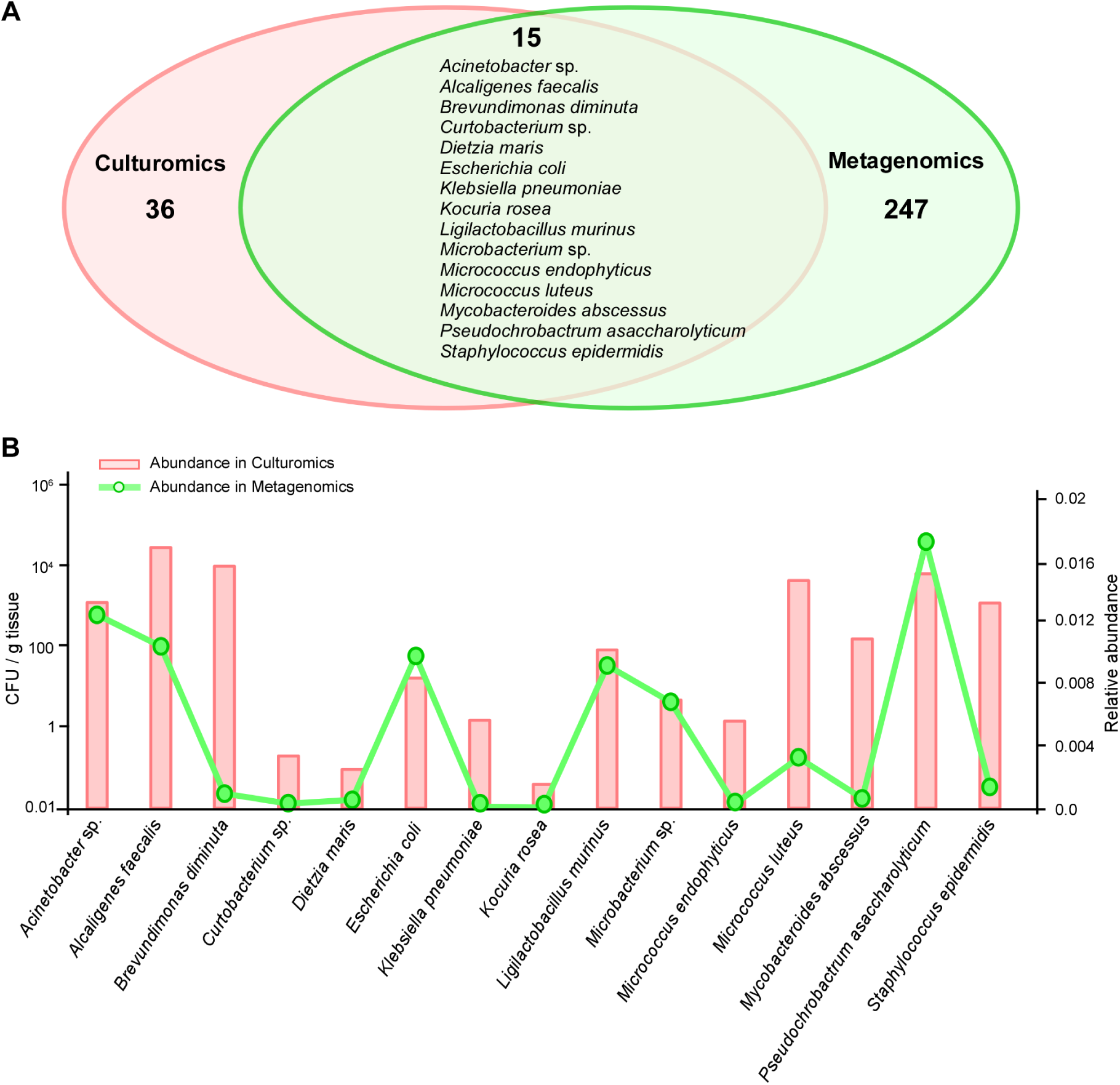
Comparative analysis of the microbial species detected by culturomics and metagenomics assays. Metagenomics assays were conducted on 10 mice (C8, C12, C13, C16, C20, C25, C31, C34, B7, and B9) that exhibited relatively high microbial abundance in the brain, heart, kidney, liver, lung, or spleen tissue. (A) Venn diagram showing microbial species detected by culturomics and metagenomics assays in the same 10 mice. The overlapping and distinct microbial species are listed in **Source Data file**. (B) Abundance of 15 common species in the 10 mice with high microbial abundance detected by culturomics (red column, CFU/g tissue) and metagenomics (green curve, relative abundance). The CFU values for culturomics assays were calculated as described in Figure 4, and the relative abundance for metagenomics assays was based on the average read coverage of 2bRAD markers. C, C57BL/6J; B, BALB/c.

**Figure 6.**
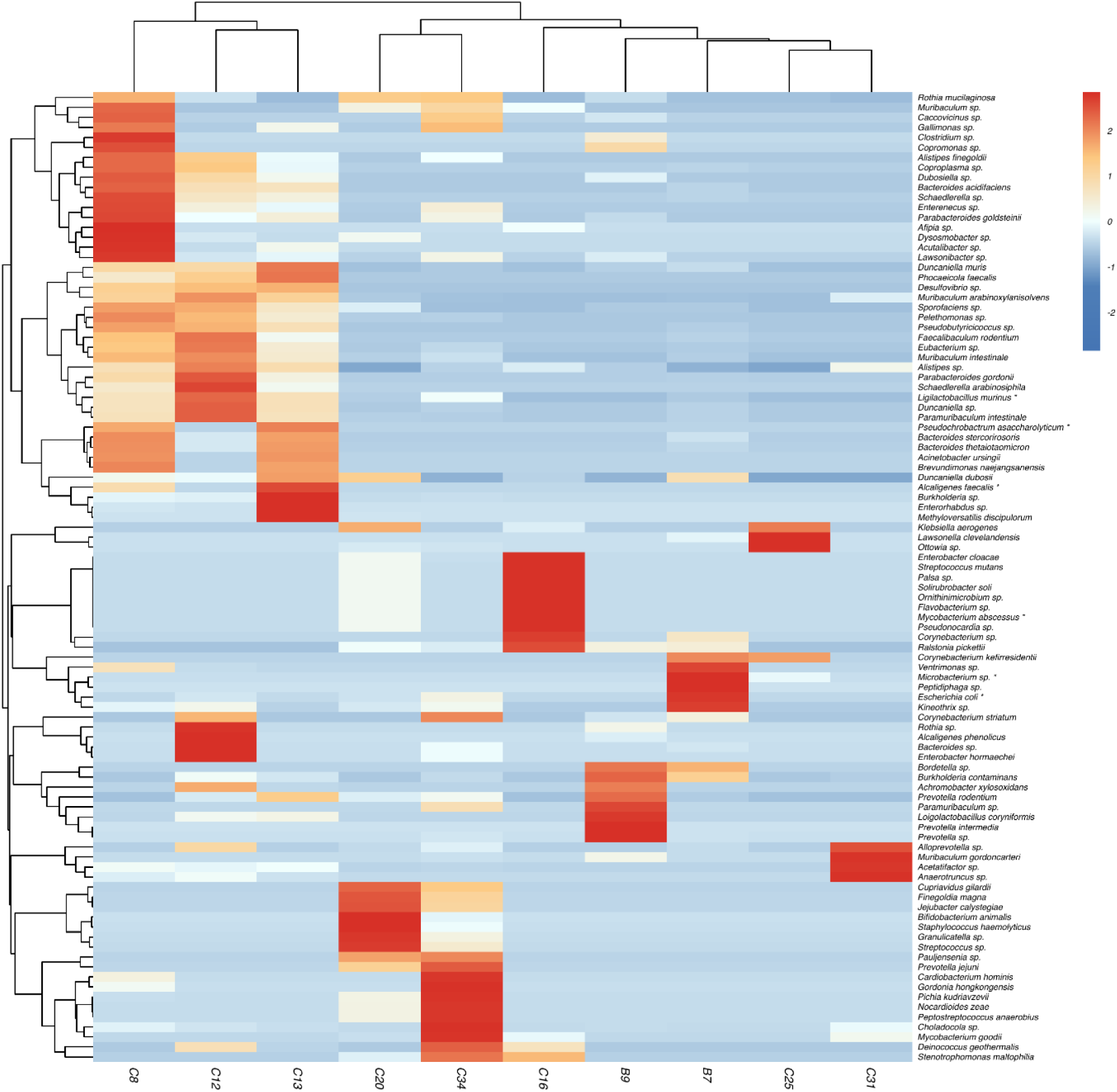
Composition and abundance of microbial species in the 10 mice with high microbial abundance detected by metagenomics assays. This analysis includes mice C8, C12, C13, C16, C20, C25, C31, C34, B7, and B9. The heatmap shows the composition and abundance of microbial species with a relative abundance >10^-4^ detected in at least two mice by metagenomics assays. The relative abundance of microbial species across brain, heart, kidney, liver, lung, and spleen tissues of each mouse was based on the average read coverage of 2bRAD markers. The intensity of red coloration indicates the relative microbial abundance, and the dendrogram shows the phylogenetic relationships of the microbial cohorts. The * symbol marks species identified by both culturomics and metagenomics assays. Detailed information for the mice, microbial species, and relative abundance is provided in **Dataset S4**.

We next investigated whether the microbial species distribution exhibited organ specificity based on the metagenomics data. As shown in **Figure S5A and Dataset S5**, distinct microbiome features were observed across different mouse organs, with specific phyla predominating in each organ. For example, *Bacteroidota*, *Pseudomonadota*, and *Bacillota* were predominant in the liver, while *Bacillota* was the most abundant phylum in the lung. The kidney and brain microbiomes were enriched in *Bacteroidota*, and *Bacillota*, while the heart microbiome exhibited a higher burden of *Acidobacterioa*, *Actinomycetota*, and *Bacteroidota*.

At the genus level, species distributions also exhibited organ-specific patterns (**Figure S5B and Dataset S5**). For example, *Alcaligenes*, *Rothia*, *Brevundimonas*, and *Pseudochrobactrum* were enriched in the brain, while *Duncaniella, Acinetobacter*, and *Mycobacterium* were predominant in the heart. In the kidney, *Rothia*, *Ligilactobacillus*, and *Alistipes* were common, while *Ligilactobacillus*, *Alistipes*, *Sphingomonas*, and *Bordetella* were enriched in the liver. In the lung, *Alistipes*, *Streptococcus,* and *Burkholderia* were dominant, and in the spleen, *Pseudomonas*, *Corynebacterium*, and *Alcaligenes* were the most prevalent genera. Consistently, *Ligilactobacillus* species were also enriched in the kidney and liver, while *Alcaligenes* and *Acinetobacter* species were identified in the brain and heart by culturomics assays.

At the species level, distinct dominant species were identified in different organs (**Dataset S5**). For example, *Muribaculum arabinoxylanisolvens, Alistipes* sp., and *Duncaniella dubosii,* were enriched in all six organs, while *L. murinus*, *Duncaniella muris*, and *Eubacterium* sp., were abundant in the brain, kidney, liver, lung, and spleen. *Muribaculum intestinale*, *Duncaniella* sp., *Paramuribaculum intestinale*, and *Anaerotruncus* sp., were abundant in four organs, while *Kineothrix* sp., *Acetatifactor* sp., and *Parabacteroides gordonii*, were enriched in three organs. *Ralstonia pickettii, Escherichia coli, Alcaligenes faecalis, Faecalibaculum rodentium* was enriched in the brain, heart, liver, and spleen, respectively. *Alistipes* sp., and *Duncaniella dubosii* were detected in more than 8 mice by metagenomics assays (**Figure 6, Dataset S4**). In conclusion, metagenomics analysis has revealed the presence of microbial species in the "sterile" organs of experimental mice that are difficult to culture. These findings suggest that species such as *Alistipes sp.*, *Duncaniella dubosii*, and others may be dominant in these organs but are not readily identified through traditional culturomics approaches.

### A potential link between the fecal and tissue microbiomes

The gut of animals harbors a diverse community of microbes that significantly impact various aspects of host health [53]. To investigate a potential link between the gut microbiome and the microbiome of “sterile” organs, we performed metagenomics analysis on the fecal microbiomes of mice. We focus on the ten mice that exhibited a relatively high microbial abundance in the brain, heart, kidney, liver, lung, and spleen tissues (**Figure 7** and **Source data**). Our analysis revealed that four common phyla (15.4%) were abundant in both fecal and tissue microbiomes. These included *Bacteroidota, Bacillota, Actinomycetota*, and *Pseudomonadota* (**Figure 7A**). Notably, *Pseudomonadota* and *Actinomycetota* species were more prevalent in the tissue microbiomes, while *Bacillota* and *Bacteroidota* species were enriched in the fecal microbiomes. Additionally, *Basidiomycota* was identified as a dominant phylum common to the microbiomes of the six organs, but not the fecal microbiomes.

**Figure 7.**
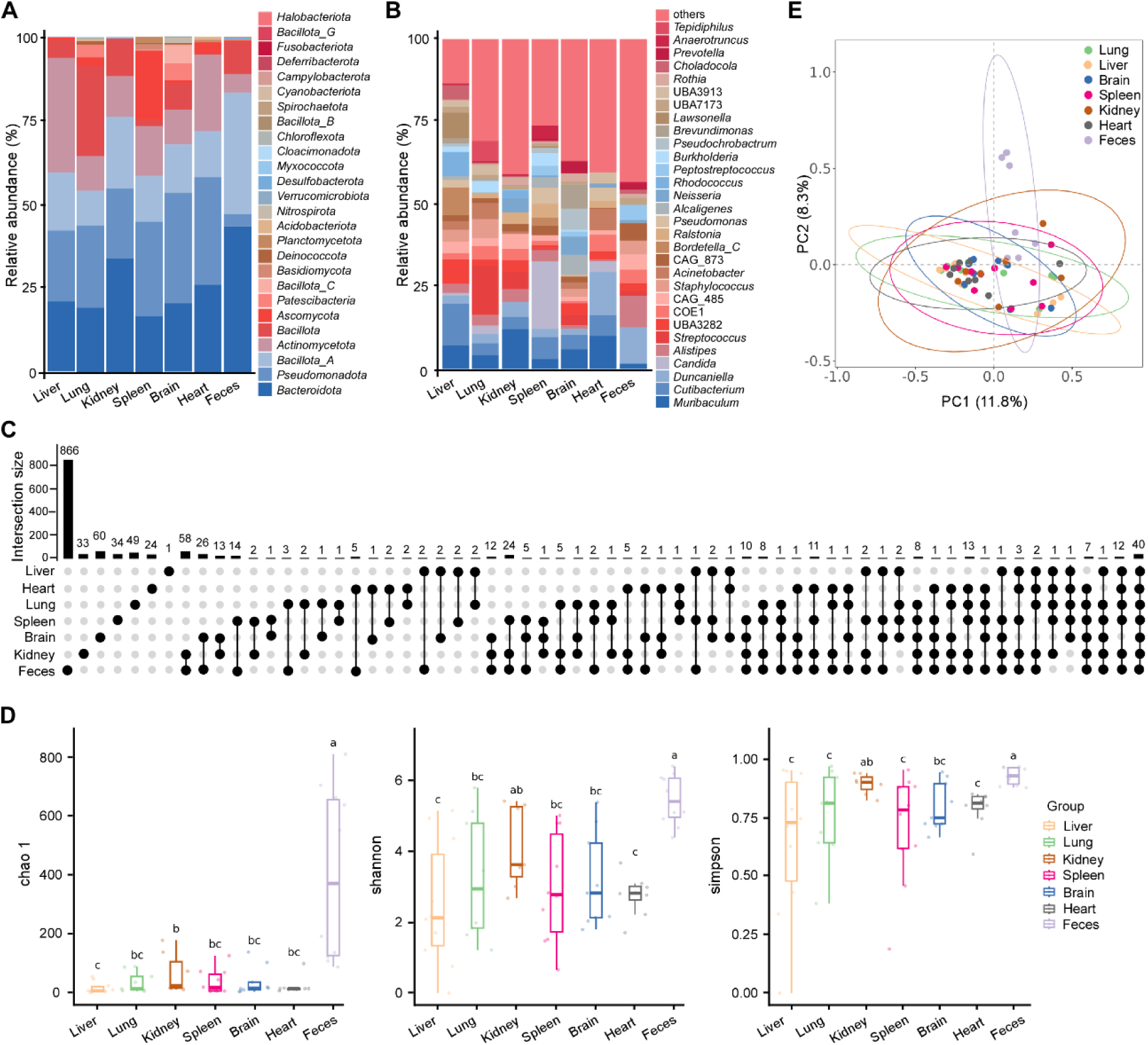
Comparative analysis of the microbiomes between the “sterile” organs and feces detected in the 10 mice with a high microbial abundance. The microbiomes in the brain, heart, kidney, liver, lung, spleen tissue, and feces of each mouse were comparatively analyzed using the 2bRAD-M method. The taxonomic composition of microbiota in different organs and feces was assigned to the phylum and genus levels based on the average relative abundance. (A and B) Comparative analysis of the dominant species detected in the organs and feces at the phylum (A) and genus (B) levels. (C) UpSet plots showing the number of common microbial species between the organ and fecal microbiomes. Columns represent different combinations of "sterile" organs and feces. (D) The organ and fecal microbiomes in healthy mice were analyzed for α-diversity measures, including Chao1, Shannon, and Simpson indices. **P<0.01. (E) Weighted PCoA plots based on Binary_Jaccard dissimilarity matrix. Each symbol represents a sample from the brain (blue), heart (gray), kidney (brown), liver (yellow), lung (green), spleen (pink), or feces (purple). The x and y axes indicate percentages of variation, and ellipses indicate the 95% confidence interval (CI). Mice detected included C8, C12, C13, C16, C20, C25, C31, C34, B7, and B9. The detailed information for the shared or specific species between the organ and fecal microbiomes is presented in **Source Data file**.

At the genus level, we analyzed the top 30 abundant taxa in the fecal microbiomes, as shown in **Figure 7B**. Of these, 10 genera (33.3%) were also detected in the microbiomes of the six organs. These included *Muribaculum*, *Duncaniella*, *Alistipes*, *Ralstonia*, *Rothia*, *Choladocola*, *Acetatifactor*, *Paramuribaculum*, *Pelethomonas*, and *Faecalibaculum*. Among these, *Muribaculum* and *Ralstomia* were more abundant in the tissue microbiomes, while *Alistipes*, *Faecalibaculum*, *Pelethomonas*, and *Paramuribaculum* were more abundant in the fecal microbiomes. Notably, *Cutibacterium* species were specifically enriched in the organ microbiomes (**Figure 7B**). We then performed an UpSet plot analysis to explore the overlap of microbial species between the organ and fecal microbiomes (**Figure 7C**). The fecal microbiomes showed the highest diversity, while the liver tissue microbiome had the lowest diversity. Tissue-specific microbial species were also identified in the six “sterile” organs (**Figure 7C**). Forty microbial species were found in both the organ and fecal microbiomes. Among these, *L. murinus* was most frequently identified in the organs using both culturomics and metagenomics assays. Since *L. murinus* is highly abundant in the fecal microbiomes, this suggests a potential translocation of microbial cells from the gut to the “sterile” organs. Furthermore, nine species, including *Alistipes* sp., *Anaerotruncus* sp., *Duncaniella dubosii*, *Duncaniella muris*, *Eubacterium* sp., *L. murinus*, *Muribaculum arabinoxylanisolvens*, *Muribaculum intestinale*, and *Paramuribaculum intestinale*, were dominant in both the organs and feces. Alpha diversity measures indicated a significantly lower richness of species in the organ microbiomes compared to the feces (**Figure 7D**). Weighted principal coordinate analysis (PCoA) confirmed distinct differences in microbial composition between the organ and fecal microbiomes (**Figure 7E**). These results suggest that, compared to the fecal microbiomes, the microbial profiles of the “sterile” organs exhibit more similarity. Both common and different microbial species were identified in the organ and fecal microbiomes, suggesting that the organs may represent unique ecological niches. Moreover, since most of the intratissue microbial species could also be found in the fecal microbiota, indicating a potential link to the gut microbiota.

### Detection of *Ligilactobacillus murinus* in the “sterile” organs and mesenteric lymph nodes (MLNs)

Since *L. murinus* was one of the most frequently isolated bacterium from the mouse “sterile” organs (**Figures 8 and S6**), we next re-evaluated the abundance of microbial organisms in these tissues and MLNs using MRS medium (an optimal medium for the growth of *L. murinus*). Ten C57BL/6J mice purchased from ZYU were analyzed as described earlier. As shown in **Figure 8**, *L. murinus* was detected in MLNs of all the 10 mice, in the brain and lung tissues of two mice, in the kidney and spleen of four mice, and in the liver of five mice. In eight out of ten mice, the abundance of *L. murinus* in the MLNs was higher than that in the other “sterile” organs. In the heart tissue, the bacterial burdens were extremely low and no *L. murinus* cells were detected. These findings imply that *L. murinus* cells in the mouse “sterile” organs could be translocated from the gut microbiota to MLNs and then to the other remote “sterile” organs.

**Figure 8.**
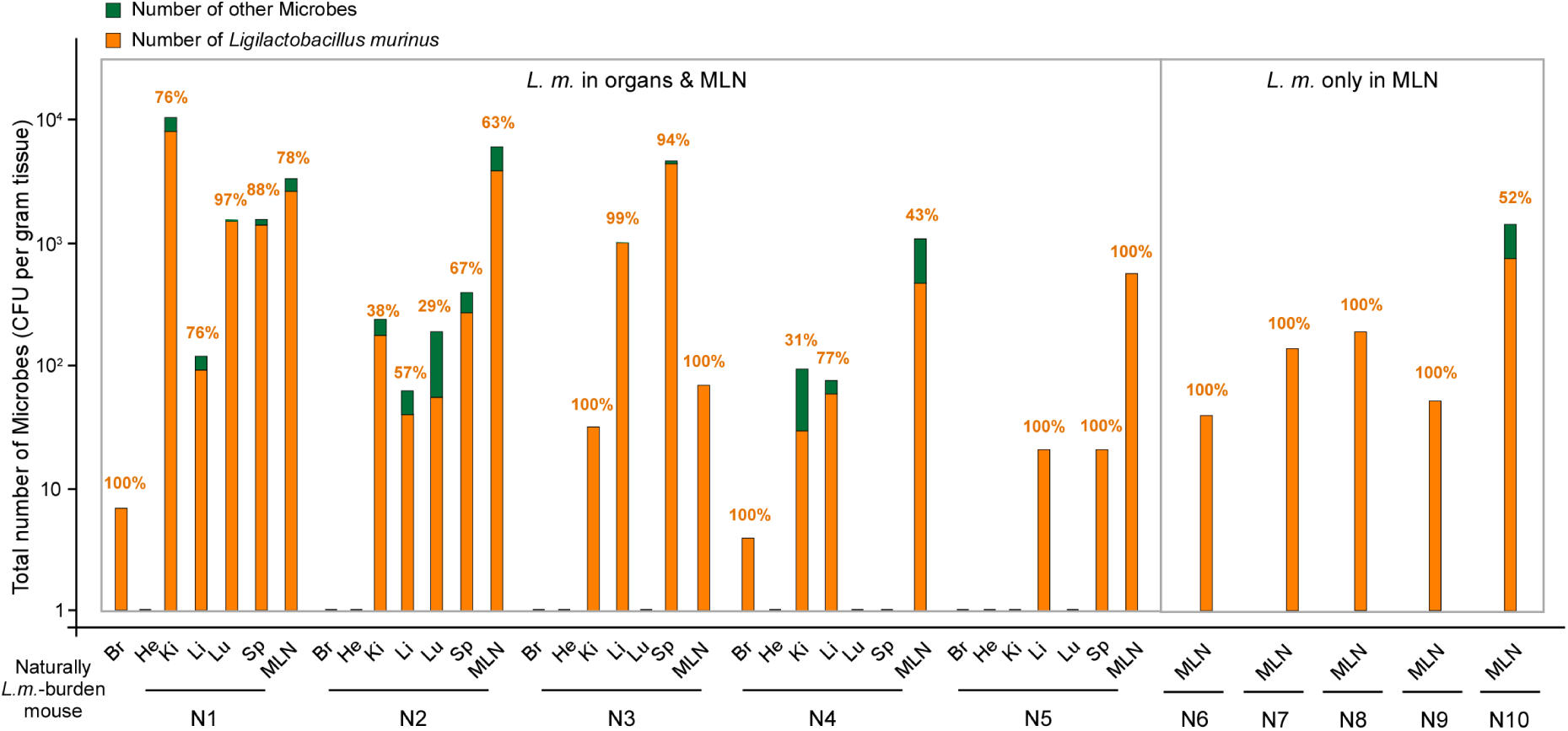
The distribution of *L. murinus* in the mouse “sterile” organs and Mesenteric Lymph Nodes (MLNs). Microbial burdens were evaluated in a total of 10 C57BL/6J mice purchased from ZYU. Upon arrival, the mice were immediately euthanized, and their organs were used for microbial burden analysis. After complete anesthesia, the body surface was sterilized three times with 75% ethanol. The microbial abundance (CFU/g tissue) in the six organs and MLN was evaluated at 37^°^C using MRS media (**Table S2**). CFU, colony-forming units. Columns represent number of microbes (CFU/g tissue) detected in the "sterile" organs. The orange columns represent the burdens of *L. murinus* (CFU/g tissue) and the green columns represent the burdens of other microbes (CFU/g tissue). The percentages on the columns indicate the proportions of *L. murinus* CFUs. Six organs include brain (Br), heart (He), kidney (Ki), liver (Li), lung (Lu), spleen (Sp). MLNs, Mesenteric Lymph Nodes. The left panel indicates mice that harbor *L. murinus* both in the “sterile” organs and in MLN. The right panel indicates mice that harbor *L. murinus* only in MLN. Detailed information for the mice, microbial species, and relative abundance is provided in **Source Data file**.

## Discussion

Experimental mice are widely used as animal models in biological and biomedical research due to their high physiological and genetic similarity to humans, small size, rapid reproductive cycle, and ease of maintenance in laboratory environments [54, 55]. Although most commonly used strains are inbred mice with very limited genetic variation, environmental and non-genetic factors, such as commensal microbes, can still contribute to phenotypic or experimental variability among individual mice. In this study, we aimed to explore whether living microorganisms are present in traditionally considered “sterile” organs of laboratory mice, and if so, how the organ microbiomes could influence the physiological and phenotypic variability of individual mice and the outcomes of research studies.

We analyzed the microbiomes of six organs (brain, heart, kidney, liver, lung, and spleen) from 104 laboratory mice using both culturomics and metagenomics assays. These mice were sourced from three major experimental animal providers in China, including different batches of C57BL/6J, BALB/c, ICR strains (**Figures 1, 3, S1 and Table S1**). We found that approximately 22% of mice (23/104) harbored a microbial burden of more than 1 x 10^4^ CFU/g in at least one tissue, and 5.77% of mice (6/104) had a microbial burden greater than 1 x 10^5^ CFU/g in at least one tissue. Some mice exhibited high microbial levels in multiple organs (**Figures 3, and S1**). Through culturomics, we identified 216 microbial species, while metagenomics analysis revealed 262 species. Despite variability in microbial diversity across organs, we identified a core group of microbial species present in multiple organs. Notably, six species, including *Acinetobacter* sp., *Alcaligenes faecalis*, *Escherichia coli*, *L. murinus*, *Microbacterium* sp., and *Pseudochrobactrum asaccharolyticum*, were the most frequently identified across both assays. Additionally, nine species, including *L. murinus*, *Alistipes*_sp., *Anaerotruncus*_sp., *Duncaniella_dubosii*, *Duncaniella_muris*, *Eubacterium* sp., *Muribaculum arabinoxylanisolvens*, *Muribaculum_intestinale*, and *Paramuribaculum_intestinale*, were found to be abundant in both the organ and fecal microbiomes. These findings suggest a potential link between the mouse organ and the gut, indicating possible translocation of gastrointestinal microbes to the traditionally "sterile" organs. This is consistent with previous studies suggesting that gut vascular barrier impairment can lead to the dissemination of intestinal bacteria and the onset of condition like cancer [56].

Environmental contamination is always of particular concern in the study of organ microbiome. Here, we have applied multiple approaches with stringent controls to avoid contaminations and to verify the presence of microbiome in the “sterile” organs, including microbial culture, metagenomic DNA sequencing, and FISH staining assays. To avoid potential contamination, strict disinfection assays were performed. The mice skin was comprehensively disinfected with 75% ethanol and sterile operations were performed according to aseptic-surgery protocols. The surface of the dissected tissues was washed carefully with 75% ethanol and then rinsed with sterile PBS three times. Sterile operations were performed according to aseptic-surgery protocols and the surface of dissected mouse tissues were also disinfected before used for microbial analysis. Notably, we found that no microbes were detected in some mouse organs, suggesting that disinfection assays were performed well and environmental contamination did not occur during dissection or culture procedures (**Figure 1C**). Moreover, several obligate anaerobic species were detected by using metagenomics assay, including the *Lachnoanaerobaculum* sp., *Peptostreptococcus anaerobius*, and *Prevotella* species. These findings further indicate the real existence of microbial species in the mouse tissues since the anaerobic species could not exist in the laboratory environment or mouse skin.

Given the widespread distribution of living microbes in the organs, it raises the question: how might these microbes affect the physiology of experimental mice? Could the organ microbiomes impact the outcomes of experiments and the reproducibility of research? Increasing evidence suggests that the presence of microbiomes in various tissues, including tumors and blood, may regulate the development, progression, and treatment of diseases [16–21]. For example, commensal bacteria such as *Streptococcus*, *Prevotella*, and *Veillonella* have been linked to lung function and respiratory health [57]. Dysbiosis in the lung microbiome has been associated with increased inflammation and exacerbation of respiratory symptoms [58–60]. Several studies have also reported the presence of microbiomes in tumors of the lung [16, 61], pancreas [17], liver [18], and breast tissues [19]. The microbial composition in tumors was found to be similar to that in normal adjacent tissues (NATs), suggesting that microbes in these tissues may proliferate rapidly within the unique tumor microenvironments (TMEs) [35]. However, it remains unclear how these tissues- or tumor-associated microbes influence disease progression and treatment outcomes. In our study, we demonstrated that the organs of laboratory mouse harbor a significant number of living microbes, which could serve as an invaluable model for studying the role of intratissue microbes in disease development and the interpretation of experimental results. The detection of wide distribution of *L. murinus* in the mouse organs and MLNs indicate a possible translocation of microbial cells from the gut to other “sterile” organs.

The C57BL/6J, BALB/c, and ICR mice used in this study are inbred animals, which are not true wild types and carry several genetic defects [62, 63]. Over a century of long-term laboratory passages may have introduced unknown genetic alterations, such as genetic shift and drift [47–49]. It remains unclear whether the intratissue microbes observed in these mice have become an integral part of their organs through long-term interactions and co-evolution between the host and its microbial inhabitants. Additionally, the immune systems of inbred mice may be more tolerant to microbial colonization in tissues. Interestingly, we did not observe significant differences in immune responses or cytokine expression, such as IL-1β, IL-6, IL-17, tumor necrosis factor α (TNF-α), and IFN-γ, between mice with or without a high burden of microbes (data not shown). This suggests that the intratissue microbiomes did not elicit a detectable immune response. Pathogenic bacteria were rarely detected in the mice studied. For example, *Staphylococcus aureus* cells were identified in low abundance in five mice (C8, C12, C20, C24 and B10), while *Klebsiella pneumoniae* was detected in a single mouse (C13). Given that *S. aureus* in a common commensal in both humans and animals, its presence in small number in mouse organs is unsurprising. These findings support the hypothesis that the majority of microbes identified in mouse organs may not be pathogens but rather harmless or potentially beneficial commensals.

## Conclusion

In summary, our findings reveal the widespread presence of living microbiomes in the traditionally considered “sterile” organs of laboratory mice with different genetic backgrounds. The use of both culturomics and metagenomics assays suggests a potential link between the organ and gastrointestinal (GI) microbiomes. However, this study also highlights several limitations. 1) Detection Limitations: Culturomics assays were unable to identify many microbes due to the unculturable nature of certain species. Metagenomics compensated for this limitation but could not differentiate between living and dead microbes. 2) Geographical Scope: All the mice were sourced from experimental animal providers in China. It remains unclear whether similar microbial profiles are present in laboratory mice from other regions or countries. 3) Dynamic Changes: We were unable to examine the dynamics of organ microbiomes within the same mice over time. Consequently, it remains uncertain whether the abundance of microbes in the organs is maintained through intratissue reproduction or continuous translocation from the GI microbiome. 4) Physiological Impact: The physiological significance of intratissue microbes remains to be explored. It is unclear how variations in microbial diversity and abundance might influence individual mouse physiology and whether these differences contribute to variability in experimental outcomes. These findings raise several intriguing questions for future research. Are living microbes also present in the "sterile" organs of other healthy experimental animals or humans? If so, do these microbes contribute to homeostasis or play a role in disease? Could certain microbial species from "sterile" tissues be harnessed for therapeutic purposes? Further investigations are necessary to determine the origin, functional roles, and potential applications of intratissue microbes, as well as to develop strategies to minimize microbiome-related variability in research outputs.

## Materials and Methods

### Animals

A total of 104 healthy mice were purchased from three major experimental animal providers in China: WTL, NMO, and ZYU. These included 44 C57BL/6J, 30 BALB/c (male, 7 weeks), and 30 ICR (male, 7 weeks) strains (**Table S1**). 44 C57BL/6J mice included 41 male mice aged 4, 6, 7 weeks, and 3, 12, 21 months, respectively, and 3 female mice aged 4 and 7 weeks. All animal experiments were conducted in compliance with the guidelines of the Animal Care and Use Committee of Fudan University (2021JS004). Ethical approval for this study was obtained from the same committee.

### Sample collection and processing

Upon acquisition, the mice were humanely euthanized. Brain, heart, liver, spleen, lung and kidney tissues were collected for microbiome analysis. The mice were first weighed and then intraperitoneally injected with 1% pentobarbital (100 mg/kg) for anesthesia. To avoid potential contamination, strict disinfection assays were performed. The mice skin was comprehensively disinfected with 75% ethanol and sterile operations were performed according to aseptic-surgery protocols. The mice were dissected, and blood samples were carefully collected from the heart ventricles. To prevent contamination, all procedures were conducted under sterile conditions in accordance with aseptic surgery protocols. The six organs (heart, lung, liver, spleen, kidney, and brain) were harvested sequentially. The surface of the dissected tissues was washed carefully with 75% ethanol and then rinsed with sterile PBS three times. Subsequently, each organ was weighed and divided into six equal portions. One portion was immediately frozen in liquid nitrogen and stored at -80 °C for metagenomic sequencing analysis. Another portion was fixed in 10% formalin (G2162, Solarbio, China) and stored at 4 °C for tissue sectioning. The remaining portions were freshly ground and processed in sterile tubes with distinct liquid media for culturomics. To minimize cross-contamination, a separate set of surgical instruments was used for each organ.

Fecal samples from each mouse were collected in sterile tubes, rapidly frozen in liquid nitrogen, and stored at -80 °C until further use.

### Culturomics

Culturomics studies were conducted following a previously described protocol with slight modifications [64]. Briefly, three tissues samples from each organ were cultured in different liquid media to target specific microbial groups: BHI medium for aerobic bacterial culture, BHI medium supplemented with 0.5 g/L L-Cysteine hydrochloride hydrate (C121800, Sigma-Aldrich, USA) for anaerobic bacterial culture, and YPD medium for fungal culture. Negative control tubes (without tissue) were prepared for each condition to rule out potential contamination. All samples were homogenized using a cryogenic high-throughput tissue grinder (SCIENTZ, Ningbo, China). The resulting homogenates were plated onto various agar media and incubated under aerobically or anaerobically conditions at 37°C (**Table S2**) [65]. Anaerobic cultures were handled entirely within anaerobic incubators (D500G, GENE SCIENCE, USA). Environmental negative controls were also included to assess laboratory-borne contamination. Based on preliminary testing of 30 culture media for culturomics, seven optimal media (BHI1, BHI2, AIA, Chocolate agar, YCFA, YPD and CDA) were selected for this study (**Table S2**). These media supported microbial growth under both aerobic and anaerobic conditions at 37°C. YPD and CDA media were used specifically for fungal isolation, while the remaining five media were used for bacterial culture.

After 5 days of incubation at 37°C, colonies were selected and streaked onto fresh media for species identification. A trained colony-picking approach was employed [66, 67], with two representative colonies of each morphology (based on colony shape, color, and size) selected from each plate. Colony-forming units (CFUs) of each microbial species within an organ were estimated based on similar colony phenotype on the same plate. All selected colonies were analyzed using matrix-assisted laser desorption/ionization time-of-flight mass spectrometry (MALDI-TOF MS) [64]. Briefly, each colony was spotted onto a polished steel MALDI MSP 96 target (Bruker Daltonics, Bremen, Germany). Subsequently, 1 μL 70% formic acid (F0507, Sigma-Aldrich, USA) and 1 μL of matrix solution [saturated α-cyano acid-4-hydroxycinnamic (28166, Sigma-Aldrich, USA) in 50% acetonitrile and 2.5% trifluoroacetic acid (900666, Sigma-Aldrich, USA)] were applied in sequence. Mass spectrometry analysis was performed using an AUTOF MS1000 system (Antu, Zhengzhou, China). For isolates that could not be accurately identified using MALDI-TOF MS, further identification was performed using amplicon sequencing analysis.

### Amplicon amplification and sequencing

Genomic DNA from each isolate was extracted using TIANamp Bacteria DNA Kit (DP302, TianGen, Beijing, China) following the manufacturer’s instructions. In brief, approximately 300 μL of glass beads (G8772, Sigma-Aldrich, USA) and 200 μL of buffer GA were added to the cell pellet and homogenized using a Mini-Beadbeater-16 (607EUR, Bio-Spec, USA). Subsequently, 20 μL of protease K solution and 220 μL of buffer GB were added in sequence, followed by incubation at 70 °C for 10 min. After two rounds of washing, the genomic DNA was isolated. The 16S rRNA gene was amplified by PCR using universal primers: 27F (5’-AGAGTTTGATCCTGGCTCAG-3’) and 1492R (5’-GGTTACCTTGTTACGACTT-3’). The resulting PCR products were sent to Sangon Biotech (Shanghai, China) for Sanger sequencing. The sequences obtained were aligned with reference type strains using the NCBI BLAST algorithm for species identification.

### Metagenomics analysis

Genomic DNA from animal organs and feces was extracted using the QIAamp Fast DNA Stool Mini Kit (51604, Qiagen, Germany) according to the manufacturer’s protocol. Briefly, the samples were suspended in 400 μL of inhibitEX buffer, along with two Steel beads (BE6638, EASYBIO, China) and ∼300 μL of glass beads (G8772, Sigma-Aldrich, USA), and homogenized using a cryogenic high-throughput tissue grinder (SCIENTZ, Ningbo, China). After centrifugation at 12000rpm for 5 min, 15 μL of protease K solution and 200 μL of buffer AL were added to the supernatant. The mixture was incubated at 70°C for 10 min, followed by the addition of 200 μL of anhydrous ethanol. The entire mixture was then transferred to an adsorption column. After two rounds of washing with buffers AW1 and AW2, genomic DNA was eluted. Negative control samples (no tissue) underwent DNA extraction simultaneously to account for potential contamination.

2bRAD-M sequencing was performed by OE Biotech (Qingdao, China) to identify microbiomes in animal tissues and feces. The 2bRAD-M library was prepared following the protocol developed by Wang *et al*. with minor modifications [68]. Briefly, genomic DNA (1 pg-200 ng) was digested with 4 U of the enzyme BcgI (NEB), and 800 U of T4 DNA ligase (NEB) was used to ligate adaptors to both ends of the DNA fragments. The ligation products were amplified, and the PCR products were resolved using 8% polyacrylamide gel. DNA bands of approximately 100 bp were excised and treated with nuclease-free water for 6-12 h at 4 °C. Sample-specific barcodes were introduced via PCR using platform-specific barcode-bearing primers. The 20 µL PCR reaction contained 6 µL of gel-extracted PCR product, 0.2 µM of each primer, 0.3 mM dNTP, 1×Phusion HF buffer, and 0.4 U of Phusion high-fidelity DNA polymerase (NEB). PCR products were purified using the QIAquick PCR purification kit (Qiagen, Germany) and sequenced on the Illumina Nova PE150 platform (Illumina, USA).

The genome information of 404,199 microbes was downloaded from the GTDB and Ensembl databases. Sixteen type 2B restriction enzymes were used to sample restriction fragments from the genomes, creating a comprehensive 2bRAD microbial genome database. The set of 2bRAD tags derived from each genome was associated with its GCF number [69]. Sequenced 2bRAD tags were mapped to the 2bRAD marker database, which together formed the 2bRAD marker database with a G-score applied to control false-positive identifications [70]. The average read coverage of 2bRAD markers per species was calculated, representing relative abundance of each species in the sample at the given sequencing depth.

### Fluorescence In Situ Hybridization (FISH)

The dissected tissues were fixed in 10% formalin and stored at 4 °C. Tissue sectioning and staining were performed by Seryicebio Inc. (Wuhan, China). Briefly, tissue blocks from the target area were cut into ∼3 mm slices, dehydrated through a gradient of alcohols (from low to high concentrations), and embedded in paraffin wax. The embedded tissues were sectioned at 10 μm intervals, with each section having a thickness of 4 μm. The sections were then deparaffinized and rehydrated sequentially. Each tissue section was hybridized with two probes simultaneously: a Cy3-labeled universal bacterial probe, EUB338 (5’-GCTGCCTCCCGTAGGAGT-3’), which fluoresced red, and a 6-FAM-labeled nonspecific complement probe (5’-CGACGGAGGGCATCCTCA-3’), which fluoresced green [56]. The nonspecific probe served as a negative control to confirm the absence of non-specific hybridization. Following hybridization, the sections were washed to remove the hybridization solution. The nuclei were counterstained with diamidino-phenyl-indole (DAPI). Finally, the sections were mounted using antifade solution (G1401, Seryicebio, Wuhan, China). Images were captured using an Ortho-Fluorescent Microscopy (NIKON ECLIPSE CI, Nikon, Japan). Fluorescence signals were observed under the following conditions: FAM (488) emitted green fluorescence with an excitation wavelength of 465-495 nm and an emission wavelength of 515-555 nm, while CY3 emitted red fluorescence with an excitation wavelength of 510-560 nm and an emission wavelength of 590 nm.

### Phylogenetic analysis

A list of 16S rRNA gene sequences was compiled from high-abundance strains identified through Culturomics. The sequences were aligned by ClustalW, and phylogenetic inference was performed using the Neighbor-Joining method in MEGA11 software. A total of 39 phylogenetic trees of high-abundance isolates were constructed. The visualization and further processing of the evolutionary trees were conducted using the iTOL platform (https://itol.embl.de/).

### Bacterial burden assay for *Ligilactobacillus murinus* using MRS medium

Ten C57BL/6J mice (7 weeks, male) purchased from ZYU were used for bacterial burden analysis. The six “sterile” organs (heart, lung, liver, spleen, kidney, and brain) and MNLs were harvested and weighted sequentially. As described earlier, the tissues were homogenized and were plated onto the MRS agar medium (an optimal medium for the growth of *L. murinus*) and incubated under aerobically conditions at 37°C (**Table S2**) [65]. The identification of bacteria was achieved using MALDI-TOF MS and 16S rRNA gene sequencing assays.

### Statistical analysis and reproducibility

Statistical analyses were performed using OECloud tools (https://cloud.oebiotech.com/) and open-source R software. α-diversity and microbial community comparisons were evaluated using ANOVA permutation and Kruskal Wallis tests. Binary_Jaccard distances were statistically analyzed using permutational multivariate analysis of variance (PERMANOVA) to assess differences in β-diversity. All statistical tests were two-sided, and results with P<0.05 were considered statistically significant.

## Data availability

The authors declare that the data supporting the findings of this study are included in the article and its Supplementary Information files. All raw sequencing data generated in this study have been deposited in the NCBI Sequence Read Archive (SRA) under accession number PRJNA1193792. Source data are provided within this paper.

## Acknowledgments

This work was supported by the National Natural Science Foundation of China (grants 31930005 and 82272359 to GH, grants 32170194 and 32370202 to LT). The content is solely the responsibility of the authors and does not represent the views of the funders. The funders had no role in the study design; data collection, analyses, or interpretation; the writing of the manuscript; or the decision to publish the results.

## Author contributions

L.T. and G.H. conceived and designed the study. M.X., S.G., C.Z. and M.M. performed the experiments and data analyses. M.X., L.T. and G.H. prepared figures, performed data interpretation and wrote/edited the manuscript. All authors read, commented, and approved the manuscript.

## References

1. Byrd AL, Belkaid Y, Segre JA. The human skin microbiome. Nat Rev Microbiol. 2018;16(3):143–55. doi: 10.1038/nrmicro.2017.157.

2. Eckburg PB, Bik EM, Bernstein CN, Purdom E, Dethlefsen L, M. S, et al. Diversity of the human intestinal microbial flora. Science. 2005;308(5728):1635-8. doi: 10.1126/science.1110591.

3. Baker JL, Mark Welch JL, Kauffman KM, McLean JS, He X. The oral microbiome: diversity, biogeography and human health. Nat Rev Microbiol. 2024;22(2):89–104. doi: 10.1038/s41579-023-00963-6.

4. Kuziel GA,Rakoff-Nahoum S. The gut microbiome. Curr Biol. 2022;32(6):R257-R64. doi: 10.1016/j.cub.2022.02.023.

5. Ma B, Forney LJ, Ravel J. Vaginal microbiome: rethinking health and disease. Annu Rev Microbiol. 2012;66(1):371–89. doi: 10.1146/annurev-micro-092611-150157.

6. Zheng D, Liwinski T, Elinav E. Interaction between microbiota and immunity in health and disease. Cell Res. 2020;30(6):492–506. doi: 10.1038/s41422-020-0332-7.

7. Hacquard S, Garrido-Oter R, González A, Spaepen S, Ackermann G, Lebeis S, et al. Microbiota and host nutrition across plant and animal kingdoms. Cell Host Microbe. 2015;17(5):603–16. doi: 10.1016/j.chom.2015.04.009.

8. Dethlefsen L, McFall-Ngai M, Relman DA. An ecological and evolutionary perspective on human-microbe mutualism and disease. Nature. 2007;449(7164):811-8. doi: 10.1038/nature06245.

9. Macpherson AJ, Geuking MB, McCoy KD. Immune responses that adapt the intestinal mucosa to commensal intestinal bacteria. Immunology. 2005;115(2):153–62. doi: 10.1111/j.1365-2567.2005.02159.x.

10. Chu H, Mazmanian SK. Innate immune recognition of the microbiota promotes host-microbial symbiosis. Nat Immunol. 2013;14(7):668–75. doi: 10.1038/ni.2635.

11. Zhang M, Sun K, Wu Y, Yang Y, Tso P, Wu Z. Interactions between intestinal microbiota and host immune response in inflammatory bowel disease. Frontiers in Immunology. 2017;8. doi: 10.3389/fimmu.2017.00942.

12. Garrett WS. Cancer and the microbiota. Science. 2015;348(6230):80–6. doi: 10.1126/science.aaa4972.

13. Mazmanian SK, Liu CH, Tzianabos AO, Kasper DL. An immunomodulatory molecule of symbiotic bacteria directs maturation of the host immune system. Cell. 2005;122(1):107–18. doi: 10.1016/j.cell.2005.05.007.

14. Morrison DJ, Preston T. Formation of short chain fatty acids by the gut microbiota and their impact on human metabolism. Gut Microbes. 2016;7(3):189–200. doi: 10.1080/19490976.2015.1134082.

15. Michán-Doña A, Vázquez-Borrego MC, Michán C. Are there any completely sterile organs or tissues in the human body? Is there any sacred place? Microb Biotechnol. 2024;17(3). doi: 10.1111/1751-7915.14442.

16. Greathouse KL, White JR, Vargas AJ, Bliskovsky VV, Beck JA, von Muhlinen N, et al. Interaction between the microbiome and TP53 in human lung cancer. Genome Biol. 2018;19(1). doi: 10.1186/s13059-018-1501-6.

17. Pushalkar S, Hundeyin M, Daley D, Zambirinis CP, Kurz E, Mishra A, et al. The pancreatic cancer microbiome promotes oncogenesis by induction of innate and adaptive immune suppression. Cancer Discovery. 2018;8(4):403–16. doi: 10.1158/2159-8290.Cd-17-1134.

18. Xue C, Jia J, Gu X, Zhou L, Lu J, Zheng Q, et al. Intratumoral bacteria interact with metabolites and genetic alterations in hepatocellular carcinoma. Signal Transduction and Targeted Therapy. 2022;7(1). doi: 10.1038/s41392-022-01159-9.

19. Urbaniak C, Gloor GB, Brackstone M, Scott L, Tangney M, Reid G, et al. The microbiota of breast tissue and its association with breast cancer. Appl Environ Microbiol. 2016;82(16):5039–48. doi: 10.1128/aem.01235-16.

20. Poore GD, Kopylova E, Zhu Q, Carpenter C, Fraraccio S, Wandro S, et al. Microbiome analyses of blood and tissues suggest cancer diagnostic approach. Nature. 2020;579(7800):567-74. doi: 10.1038/s41586-020-2095-1.

21. Apostolou P, Tsantsaridou A, Papasotiriou I, Toloudi M, Chatziioannou M, Giamouzis G. Bacterial and fungal microflora in surgically removed lung cancer samples. J Cardiothorac Surg. 2011;6(137):1–5. doi: 10.1186/1749-8090-6-137.

22. Helmink BA, Khan MAW, Hermann A, Gopalakrishnan V, Wargo JA. The microbiome, cancer, and cancer therapy. Nat Med. 2019;25(3):377–88. doi: 10.1038/s41591-019-0377-7.

23. Matson V, Fessler J, Bao R, Chongsuwat T, Zha Y, Alegre M-L, et al. The commensal microbiome is associated with anti–PD-1 efficacy in metastatic melanoma patients. Science. 2018;359(6371):104-8. doi: 10.1126/science.aao3290.

24. Parhi L, Alon-Maimon T, Sol A, Nejman D, Shhadeh A, Fainsod-Levi T, et al. Breast cancer colonization by Fusobacterium nucleatum accelerates tumor growth and metastatic progression. Nat Commun. 2020;11(1). doi: 10.1038/s41467-020-16967-2.

25. Iida N, Dzutsev A, Stewart CA, Smith L, Bouladoux N, Weingarten RA, et al. Commensal bacteria control cancer response to therapy by modulating the tumor microenvironment. 2013;342:967–70. doi: 10.1126/science.1240527.

26. Routy B, Chatelier EL, Derosa L, Duong CPM, Alou MT, Daillère R, et al. Gut microbiome influences efficacy of PD-1-based immunotherapy against epithelial tumors. 2018;359:91–7. doi: 10.1126/science.aan3706.

27. Gopalakrishnan V, Spencer CN, Nezi L, Reuben A, Andrews MC, Karpinets TV, et al. Gut microbiome modulates response to anti–PD-1 immunotherapy in melanoma patients. Science. 2018;359(6371):97-103. doi: 10.1126/science.aan4236.

28. Nejman D, Livyatan I, Fuks G, Gavert N, Zwang Y, Geller LT, et al. The human tumor microbiome is composed of tumor type–specific intracellular bacteria. Science. 2020;368(6494):973-80. doi: 10.1126/science.aay9189.

29. Gopalakrishnan V, Helmink BA, Spencer CN, Reuben A, Wargo JA. The influence of the gut microbiome on cancer, immunity, and cancer immunotherapy. Cancer Cell. 2018;33(4):570–80. doi: 10.1016/j.ccell.2018.03.015.

30. Wong-Rolle A, Wei HK, Zhao C, Jin C. Unexpected guests in the tumor microenvironment: microbiome in cancer. Protein Cell. 2020;12(5):426–35. doi: 10.1007/s13238-020-00813-8.

31. Farrell JJ, Zhang L, Zhou H, Chia D, Elashoff D, Akin D, et al. Variations of oral microbiota are associated with pancreatic diseases including pancreatic cancer. Gut. 2012;61(4):582–8. doi: 10.1136/gutjnl-2011-300784.

32. Sepich-Poore GD, Zitvogel L, Straussman R, Hasty J, Wargo JA, Knight R. The microbiome and human cancer. Science. 2021;371(6536). doi: 10.1126/science.abc4552.

33. Fu A, Yao B, Dong T, Chen Y, Yao J, Liu Y, et al. Tumor-resident intracellular microbiota promotes metastatic colonization in breast cancer. Cell. 2022;185(8):1356–72.e26. doi: 10.1016/j.cell.2022.02.027.

34. Leinwand JC, Paul B, Chen R, Xu F, Sierra MA, Paluru MM, et al. Intrahepatic microbes govern liver immunity by programming NKT cells. J Clin Investig. 2022;132(8). doi: 10.1172/jci158999.

35. Xie Y, Xie F, Zhou X, Zhang L, Yang B, Huang J, et al. Microbiota in tumors: from understanding to application. Adv Sci. 2022;9(21). doi: 10.1002/advs.202200470.

36. Leinwand J, Miller G. Regulation and modulation of antitumor immunity in pancreatic cancer. Nat Immunol. 2020;21(10):1152–9. doi: 10.1038/s41590-020-0761-y.

37. Riquelme E, Maitra A, McAllister F. Immunotherapy for pancreatic cancer: more than just a gut feeling. Cancer Discov. 2018;8(4):386–8. doi: 10.1158/2159-8290.Cd-18-0123.

38. Lee W-J, Hase K. Gut microbiota–generated metabolites in animal health and disease. Nat Chem Biol. 2014;10(6):416–24. doi: 10.1038/nchembio.1535.

39. Baumans V. Use of animals in experimental research: an ethical dilemma? Gene Ther. 2004;11(S1):S64–S6. doi: 10.1038/sj.gt.3302371.

40. Crabbe JC, Wahlsten D, Dudek BC. Genetics of mouse behavior interactions with laboratory environment. Science. 1999;284:1670–2. doi: 10.1126/science.284.5420.1670.

41. von Kortzfleisch VT, Karp NA, Palme R, Kaiser S, Sachser N, Richter SH. Improving reproducibility in animal research by splitting the study population into several ‘mini-experiments’. Sci Rep. 2020;10(1). doi: 10.1038/s41598-020-73503-4.

42. Leystra AA, Clapper ML. Gut microbiota influences experimental outcomes in mouse models of colorectal cancer. Genes. 2019;10(11). doi: 10.3390/genes10110900.

43. Wang X, Bing J, Zheng Q, Zhang F, Liu J, Yue H, et al. The first isolate of Candida auris in China: clinical and biological aspects. Emerg Microbes Infect. 2018;7(1):1–9. doi: 10.1038/s41426-018-0095-0.

44. Tao L, Du H, Guan G, Dai Y, Nobile CJ, Liang W, et al. Discovery of a “white-gray-opaque” tristable phenotypic switching system in Candida albicans: roles of non-genetic diversity in host adaptation. PLoS Biol. 2014;12(4). doi: 10.1371/journal.pbio.1001830.

45. Charlson ES, Bittinger K, Haas AR, Fitzgerald AS, Frank I, Yadav A, et al. Topographical continuity of bacterial populations in the healthy human respiratory tract. American Journal of Respiratory and Critical Care Medicine. 2011;184(8):957–63. doi: 10.1164/rccm.201104-0655OC.

46. Sun X, Zhang H, Zhang X, Gao W, Zhou C, Kou X, et al. The cellular microbiome of visceral organs: an inherent inhabitant of parenchymal cells. Microorganisms. 2024;12(7). doi: 10.3390/microorganisms12071333.

47. Dumont BL, Yoder A. Significant strain variation in the mutation spectra of inbred laboratory mice. Mol Biol Evol. 2019;36(5):865–74. doi: 10.1093/molbev/msz026.

48. Åhlgren J, Voikar V. Experiments done in Black-6 mice: what does it mean? Lab Anim. 2019;48(6):171–80. doi: 10.1038/s41684-019-0288-8.

49. Uchimura A, Higuchi M, Minakuchi Y, Ohno M, Toyoda A, Fujiyama A, et al. Germline mutation rates and the long-term phenotypic effects of mutation accumulation in wild-type laboratory mice and mutator mice. Genome Res. 2015;25(8):1125–34. doi: 10.1101/gr.186148.114.

50. Freeman HC, Hugill A, Dear NT, Ashcroft FM, Cox RD. Deletion of nicotinamide nucleotide transhydrogenase. Diabetes. 2006;55(7):2153–6. doi: 10.2337/db06-0358.

51. Nakamura H. BALB/c mouse. Brenner’s Encyclopedia of Genetics2013. p. 290-2.

52. Berg RD. Bacterial translocation from the gastrointestinal tract. Trends Microbiol. 1995;3(4):149–54. doi: 10.1007/978-1-4615-4143-1_2.

53. Round JL, Mazmanian SK. The gut microbiota shapes intestinal immune responses during health and disease. Nat Rev Immunol. 2009;9(5):313–23. doi: 10.1038/nri2515.

54. García-García MJ. A history of mouse genetics from fancy mice to mutations in every gene. Adv Exp Med Biol. 2020;1236:1–38. doi: 10.1007/978-981-15-2389-2_1.

55. Tam WY, Cheung K-K. Phenotypic characteristics of commonly used inbred mouse strains. J Mol Med. 2020;98(9):1215–34. doi: 10.1007/s00109-020-01953-4.

56. Bertocchi A, Carloni S, Ravenda PS, Bertalot G, Spadoni I, Lo Cascio A, et al. Gut vascular barrier impairment leads to intestinal bacteria dissemination and colorectal cancer metastasis to liver. Cancer Cell. 2021;39(5):708–24.e11. doi: 10.1016/j.ccell.2021.03.004.

57. Segal LN, Alekseyenko AV, Clemente JC, Kulkarni R, Wu B, Chen H, et al. Enrichment of lung microbiome with supraglottic taxa is associated with increased pulmonary inflammation. Microbiome. 2013;1(19). doi: 10.1186/2049-2618-1-19.

58. Ghebre MA, Pang PH, Diver S, Desai D, Bafadhel M, Haldar K, et al. Biological exacerbation clusters demonstrate asthma and chronic obstructive pulmonary disease overlap with distinct mediator and microbiome profiles. J Allergy Clin Immunol. 2018;141(6):2027–36.e12. doi: 10.1016/j.jaci.2018.04.013.

59. Haldar K, George L, Wang Z, Mistry V, Ramsheh MY, Free RC, et al. The sputum microbiome is distinct between COPD and health, independent of smoking history. Respir Res. 2020;21(1). doi: 10.1186/s12931-020-01448-3.

60. Li R, Li J, Zhou X. Lung microbiome: new insights into the pathogenesis of respiratory diseases. Signal Transduct Target Ther. 2024;9(1). doi: 10.1038/s41392-023-01722-y.

61. Panagiotis Apostolou AT, Ioannis Papasotiriou, Maria Toloudi, Marina Chatziioannou, Gregory Giamouzis. Bacterial and fungal microflora in surgically removed lung cancer samples. 2011;6:137.

62. Mekada K, Yoshiki A. Substrains matter in phenotyping of C57BL/6 mice. Exp Anim. 2021;70(2):145–60. doi: 10.1538/expanim.20-0158.

63. Mekada K, Abe K, Murakami A, Nakamura S, Nakata H, Moriwaki K, et al. Genetic differences among c57BL/6 substrains. Exp Anim. 2009;58(2):141–9. doi: 10.1538/expanim.58.141. .

64. Lagier JC, Armougom F, Million M, Hugon P, Pagnier I, Robert C, et al. Microbial culturomics: paradigm shift in the human gut microbiome study. Clin Microbiol Infect. 2012;18(12):1185–93. doi: 10.1111/1469-0691.12023.

65. Lagier J-C, Khelaifia S, Alou MT, Ndongo S, Dione N, Hugon P, et al. Culture of previously uncultured members of the human gut microbiota by culturomics. Nat Microbiol. 2016;1(12). doi: 10.1038/nmicrobiol.2016.203.

66. Chang Y, Hou F, Pan Z, Huang Z, Han N, Bin L, et al. Optimization of culturomics strategy in human fecal samples. Front Microbiol. 2019;10. doi: 10.3389/fmicb.2019.02891.

67. Alou MT, Naud S, Khelaifia S, Bonnet M, Lagier J-C, Raoult D. State of the art in the culture of the human microbiota new interests and strategies. Clin Microbiol Rev. 2021;34(1):e00129–19. doi: 10.1128/cmr.00129-19.

68. Wang S, Meyer E, McKay JK, Matz MV. 2b-RAD: a simple and flexible method for genome-wide genotyping. Nat Methods. 2012;9(8):808–10. doi: 10.1038/nmeth.2023.

69. Sun Z, Liu J, Zhang M, Wang T, Huang S, Weiss ST, et al. Removal of false positives in metagenomics-based taxonomy profiling via targeting Type IIB restriction sites. Nature Communications. 2023;14(1). doi: 10.1038/s41467-023-41099-8.

70. Sun Z, Huang S, Zhu P, Tzehau L, Zhao H, Lv J, et al. Species-resolved sequencing of low-biomass or degraded microbiomes using 2bRAD-M. Genome Biol. 2022;23(1). doi: 10.1186/s13059-021-02576-9.

