## Supplemental data for "Characterizing the Microbiome of ’Sterile’ Organs in Experimental Mice"

**This file includes:**

Figures S1 to S6

Tables S1 to S2

### Supplemental Information

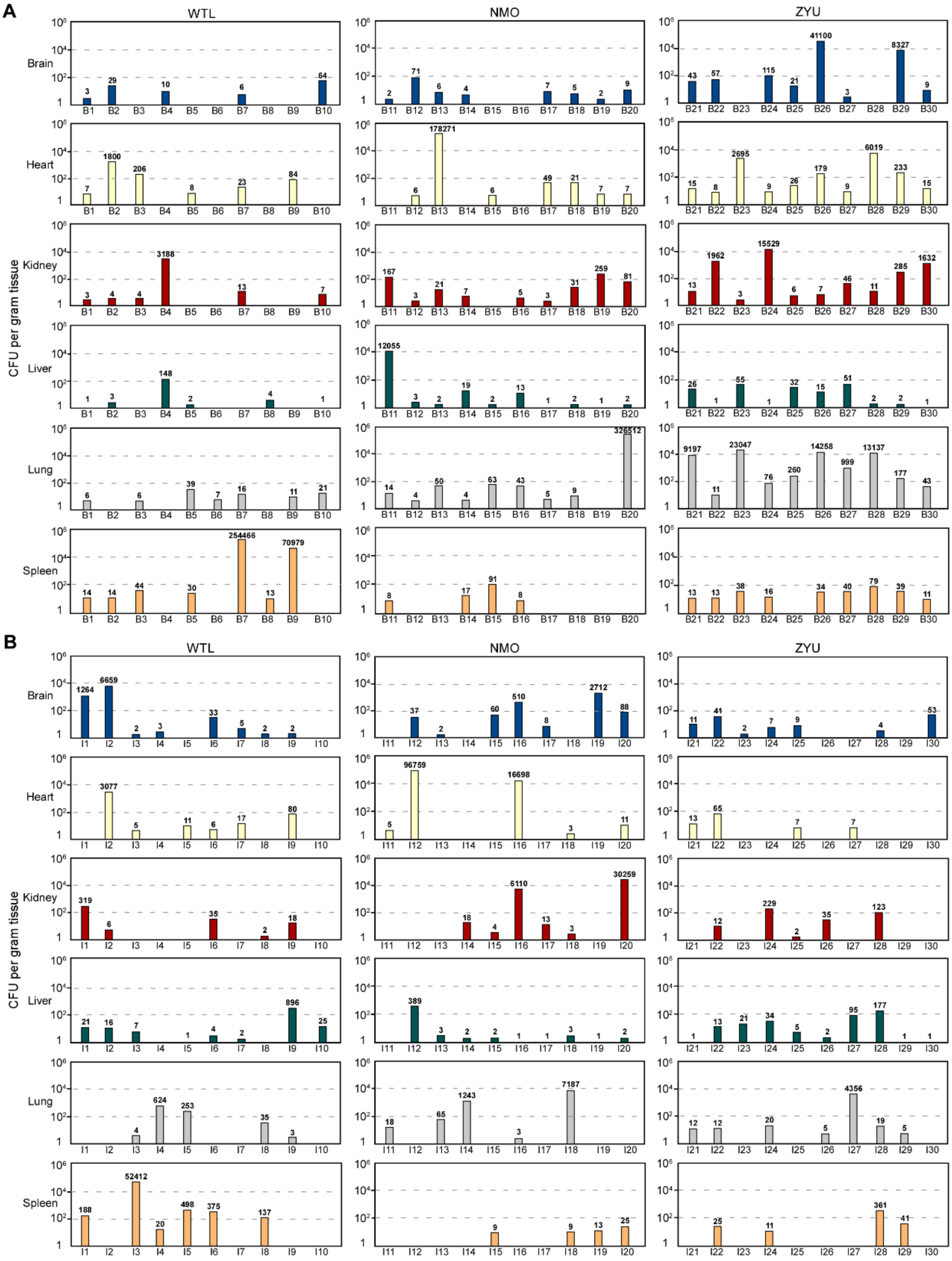


**Figure S1. Abundance of microbes detected in the brain, heart, kidney, liver, lung, and spleen tissues of BALB/c (A) and ICR (B) mice based on culturomics assays.** A total of 60 mice (30 BALB/c and 30 ICR) were purchased from three experimental animal providers in China (WTL, NMO, and

ZYU) and examined. The microbial abundance (CFU/g tissue) of the six organs of each mouse was evaluated at 37°C using the 7 optimal cultural media. The column color represents a specific organ, and the numbers on the columns indicate the CFU values. Detailed methods are described in **Figure 3**. The mouse details are shown in **Table S1**. Mice used: B1-B30 and I1-I30, mouse codes ranked by treatment order. B: BALB/c; I: ICR.

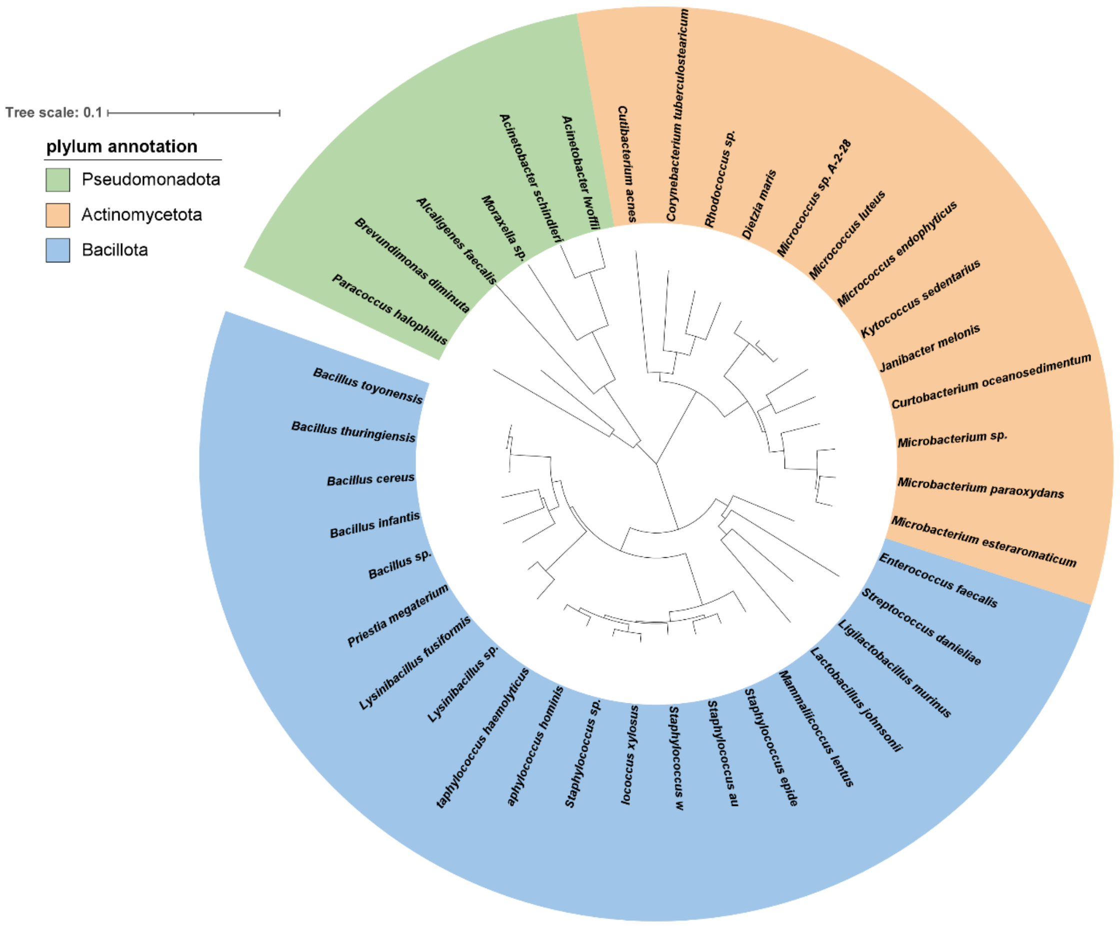


**Figure S2. Phylogenetic tree of the representative species enriched in the mouse brain, heart, kidney, liver, lung, and spleen tissues.** The species of *Pseudomonadota, Actinomycetota*, and *Bacillota* are highlighted in green, orange, and blue, respectively. The absolute total branch length is indicated at the bottom. The phylogenetic tree was based on data from MEGA11 and visualized using the iTOL platform (https://itol.embl.de/). 16S rDNA sequences of the representative species are shown in **Dataset S2**.

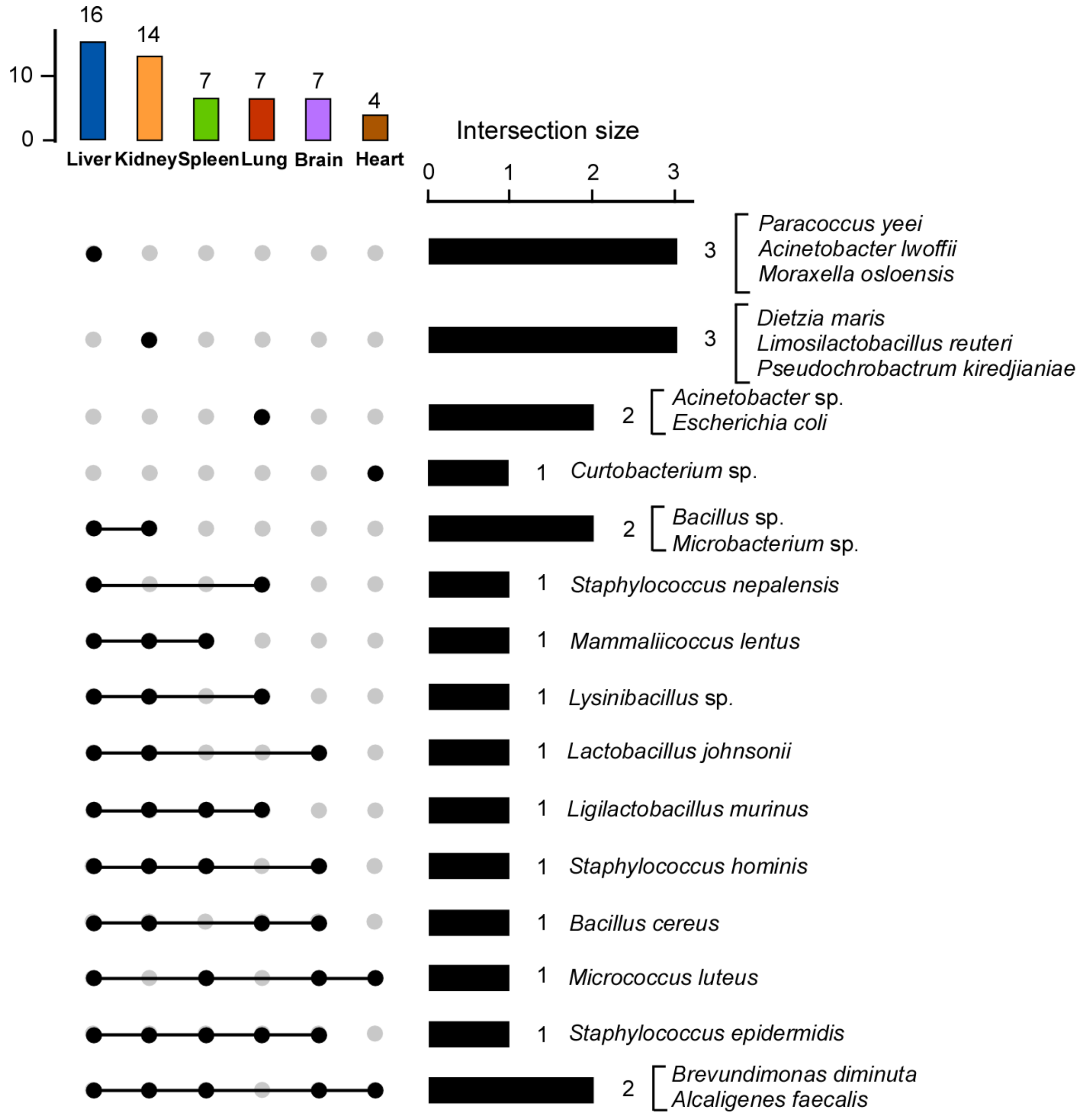


**Figure S3. Unique and overlap species among different organs of the 42 mice with a relatively high microbial burden.** The mice analyzed in this figure were shown in **Dataset S3**. 42 mice with a microbial burden higher than 1 x $10^3$ CFU/g tissues in at least one organ were detected. UpSet plots show the number of common microbial species among the six organs. Rows represent different combinations of organs. The shared or specific species were displayed on right side of the plot. The detailed information for the shared or specific species among the organs is presented in **Dataset S3**.

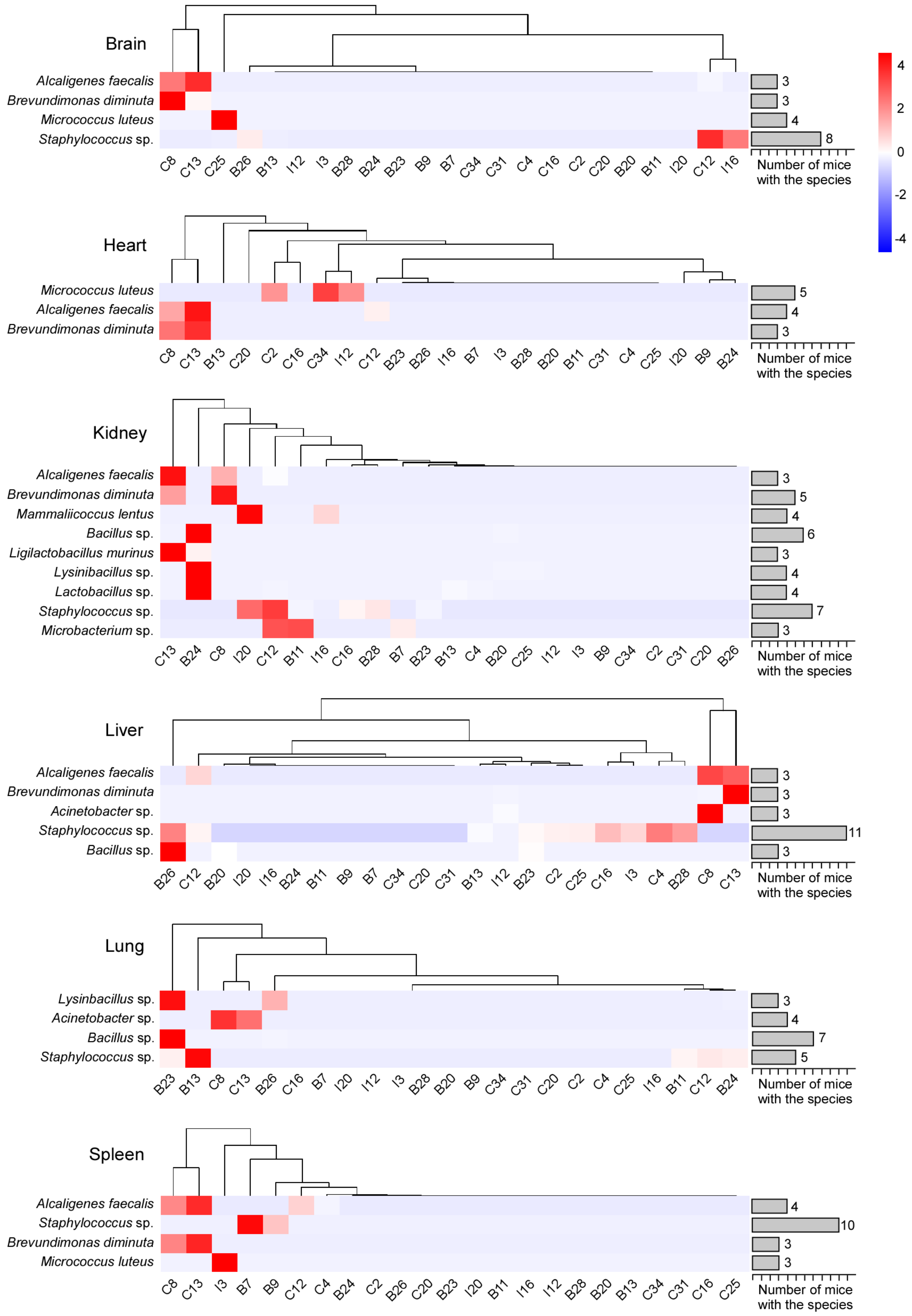


**Figure S4. Most frequently isolated microbial species from different organs of the 23 mice with a high microbial burden (>$10^4$ CFU/g tissues in at least one organ).** The mice analyzed in this figure include C2, C4, C8, C12,

C13, C16, C20, C25, C31, C34, B7, B9, B11, B13, B20, B23, B24, B26, B28, I3, I12, I16, and I20. The heatmap shows the composition and abundance of microbial species isolated from the brain, heart, kidney, liver, lung, and spleen tissues of at least three mice with a high microbial burden. Microbial abundance (CFU/g tissue) was evaluated for each organ of the individual mice. The intensity of red coloration indicates the relative microbial abundance. The dendrogram shows the phylogenetic relationships of the microbial cohorts. Number of mice harboring the corresponding species was displayed on right side of the plots. The relative microbial abundance was calculated based on the CFU of each microbial species in the corresponding organ. The heatmap was generated using OECloud tools at https://cloud.oebiotech.com. This figure is associated with **Figures 3**, **4** and **S1**. The detailed information for the mice and microbial species, as well as CFUs, is provided in **Datasets S1** and **S3**.

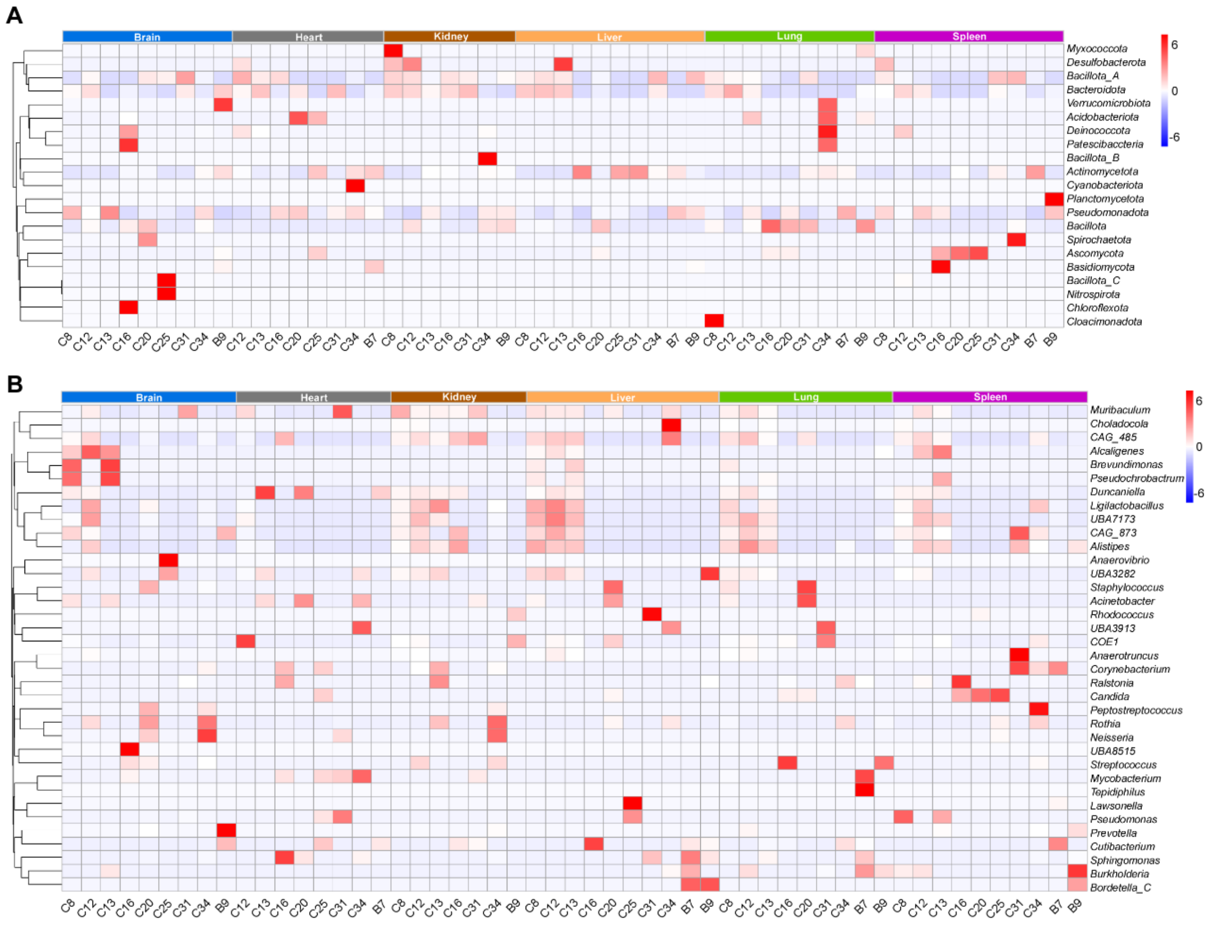


**Figure S5. Taxonomic composition of microbes in each organ of the 10 mice with a high microbial abundance detected by metagenomics assays.** Mice analyzed in this figure include C8, C12, C13, C16, C20, C25, C31, C34, B7, and B9. The relative abundance of microbial species was calculated as described in **Figure 6**. The intensity of red coloration indicates the relative microbial abundance, with darker red representing higher abundance. The dendrogram shows the phylogenetic relationships of the microbial cohorts. Organ information is indicated at the top of the images. (A) Taxonomic composition at the phylum level. (B) Taxonomic composition at the genus level. Detailed information for the mouse strains, microbial species, and relative abundance is provided in **Dataset S5**.

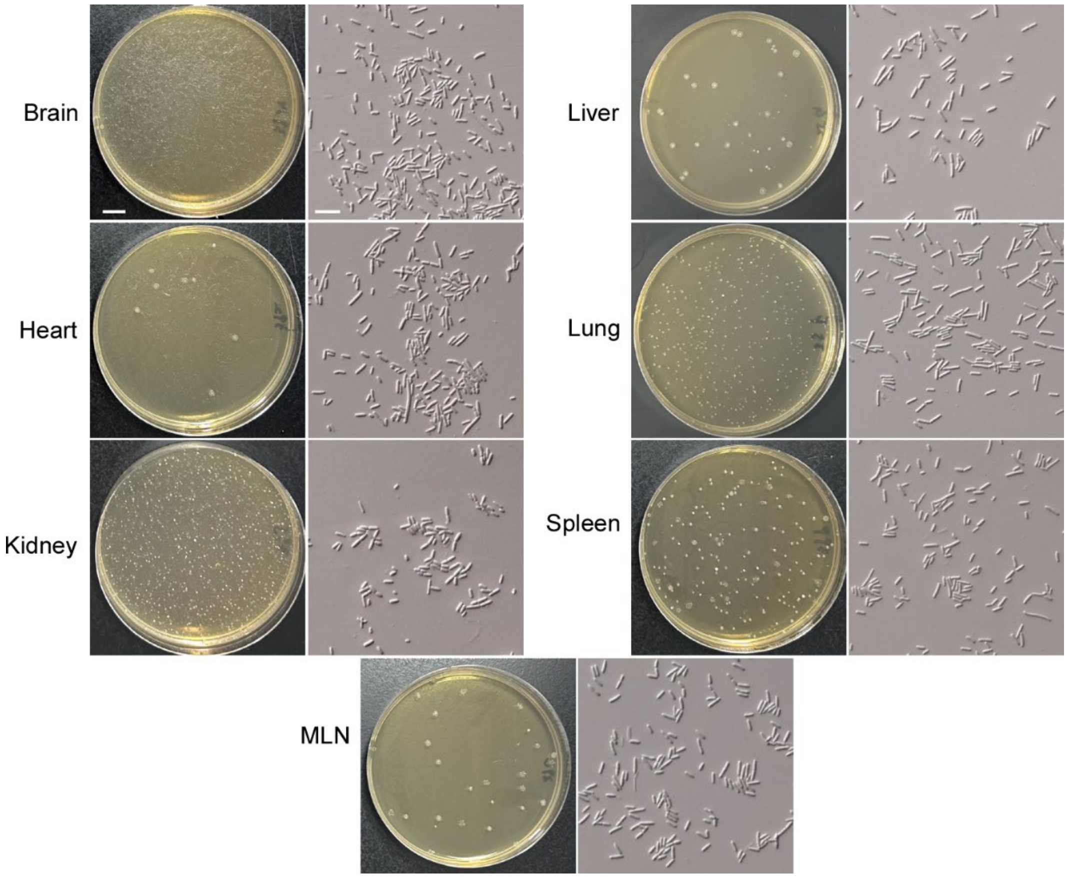


**Figure S6. Representative examples of culture phenotype for *Ligilactobacillus murinus* detection in the organs of the naturally microbe-burden mice.**

Colonial and cellular images of *Ligilactobacillus murinus* isolates derived from different organs of the naturally microbe-burden mice. MRS medium was used. 10 naturally microbe-burden mice (C57BL/6J, 7 weeks, male, n=10) were disinfected, euthanized, and dissected. Six organs including brain, heart, kidney, liver, lung, and spleen, and MLN of each mouse were harvested, weighed, and homogenized for microbial isolation. The plates were cultured under aerobic condition at 37 $^{o}$C for 5 days. Scale bar for colonies, 1 cm; Scale bar for cells, 20 μm.

**Table S1. Detailed information of the mice used in this study.**

| Mouse Code | Strain | Age | Sex | Animal provider |
|---|---|---|---|---|
| C1 | C57BL/6J | 7w | Male | WTL |
| C2 | C57BL/6J | 7w | Male | WTL |
| C3 | C57BL/6J | 7w | Male | WTL |
| C4 | C57BL/6J | 7w | Female | WTL |
| C5 | C57BL/6J | 7w | Female | WTL |
| C6 | C57BL/6J | 3m | Male | WTL |
| C7 | C57BL/6J | 3m | Male | WTL |
| C8 | C57BL/6J | 3m | Male | WTL |
| C9 | C57BL/6J | 21m | Male | WTL |
| C10 | C57BL/6J | 21m | Male | WTL |
| C11 | C57BL/6J | 21m | Male | WTL |
| C12 | C57BL/6J | 12m | Male | WTL |
| C13 | C57BL/6J | 12m | Male | WTL |
| C14 | C57BL/6J | 12m | Male | WTL |
| C15 | C57BL/6J | 4w | Female | WTL |
| C16 | C57BL/6J | 4w | Male | WTL |
| C17 | C57BL/6J | 4w | Male | WTL |
| C18 | C57BL/6J | 4w | Male | WTL |
| C19 | C57BL/6J | 4w | Male | WTL |
| C20 | C57BL/6J | 4w | Male | WTL |
| C21 | C57BL/6J | 12m | Male | WTL |
| C22 | C57BL/6J | 12m | Male | WTL |
| C23 | C57BL/6J | 12m | Male | WTL |
| C24 | C57BL/6J | 12m | Male | WTL |
| C25 | C57BL/6J | 6w | Male | NMO |
| C26 | C57BL/6J | 6w | Male | NMO |
| C27 | C57BL/6J | 6w | Male | NMO |
| C28 | C57BL/6J | 6w | Male | NMO |
| C29 | C57BL/6J | 6w | Male | NMO |
| C30 | C57BL/6J | 6w | Male | NMO |
| C31 | C57BL/6J | 6w | Male | NMO |
| C32 | C57BL/6J | 6w | Male | NMO |
| C33 | C57BL/6J | 6w | Male | NMO |
| C34 | C57BL/6J | 6w | Male | NMO |
| C35 | C57BL/6J | 7w | Male | ZYU |
| C36 | C57BL/6J | 7w | Male | ZYU |
| C37 | C57BL/6J | 7w | Male | ZYU |
| C38 | C57BL/6J | 7w | Male | ZYU |
| C39 | C57BL/6J | 7w | Male | ZYU |
| C40 | C57BL/6J | 7w | Male | ZYU |

| | | | | |
|---|---|---|---|---|
| C41 | C57BL/6J | 7w | Male | ZYU |
| C42 | C57BL/6J | 7w | Male | ZYU |
| C43 | C57BL/6J | 7w | Male | ZYU |
| C44 | C57BL/6J | 7w | Male | ZYU |
| B1 | BALB/c | 7w | Male | WTL |
| B2 | BALB/c | 7w | Male | WTL |
| B3 | BALB/c | 7w | Male | WTL |
| B4 | BALB/c | 7w | Male | WTL |
| B5 | BALB/c | 7w | Male | WTL |
| B6 | BALB/c | 7w | Male | WTL |
| B7 | BALB/c | 7w | Male | WTL |
| B8 | BALB/c | 7w | Male | WTL |
| B9 | BALB/c | 7w | Male | WTL |
| B10 | BALB/c | 7w | Male | WTL |
| B11 | BALB/c | 7w | Male | NMO |
| B12 | BALB/c | 7w | Male | NMO |
| B13 | BALB/c | 7w | Male | NMO |
| B14 | BALB/c | 7w | Male | NMO |
| B15 | BALB/c | 7w | Male | NMO |
| B16 | BALB/c | 7w | Male | NMO |
| B17 | BALB/c | 7w | Male | NMO |
| B18 | BALB/c | 7w | Male | NMO |
| B19 | BALB/c | 7w | Male | NMO |
| B20 | BALB/c | 7w | Male | NMO |
| B21 | BALB/c | 7w | Male | ZYU |
| B22 | BALB/c | 7w | Male | ZYU |
| B23 | BALB/c | 7w | Male | ZYU |
| B24 | BALB/c | 7w | Male | ZYU |
| B25 | BALB/c | 7w | Male | ZYU |
| B26 | BALB/c | 7w | Male | ZYU |
| B27 | BALB/c | 7w | Male | ZYU |
| B28 | BALB/c | 7w | Male | ZYU |
| B29 | BALB/c | 7w | Male | ZYU |
| B30 | BALB/c | 7w | Male | ZYU |
| I1 | ICR | 7w | Male | WTL |
| I2 | ICR | 7w | Male | WTL |
| I3 | ICR | 7w | Male | WTL |
| I4 | ICR | 7w | Male | WTL |
| I5 | ICR | 7w | Male | WTL |
| I6 | ICR | 7w | Male | WTL |
| I7 | ICR | 7w | Male | WTL |
| I8 | ICR | 7w | Male | WTL |
| I9 | ICR | 7w | Male | WTL |
| I10 | ICR | 7w | Male | WTL |

| | | | | |
|---|---|---|---|---|
| I11 | ICR | 7w | Male | NMO |
| I12 | ICR | 7w | Male | NMO |
| I13 | ICR | 7w | Male | NMO |
| I14 | ICR | 7w | Male | NMO |
| I15 | ICR | 7w | Male | NMO |
| I16 | ICR | 7w | Male | NMO |
| I17 | ICR | 7w | Male | NMO |
| I18 | ICR | 7w | Male | NMO |
| I19 | ICR | 7w | Male | NMO |
| I20 | ICR | 7w | Male | NMO |
| I21 | ICR | 7w | Male | ZYU |
| I22 | ICR | 7w | Male | ZYU |
| I23 | ICR | 7w | Male | ZYU |
| I24 | ICR | 7w | Male | ZYU |
| I25 | ICR | 7w | Male | ZYU |
| I26 | ICR | 7w | Male | ZYU |
| I27 | ICR | 7w | Male | ZYU |
| I28 | ICR | 7w | Male | ZYU |
| I29 | ICR | 7w | Male | ZYU |
| I30 | ICR | 7w | Male | ZYU |

**Notes:** w: week, m: month. All the mice used in this study were from three major experimental animal providers in China (WTL, NMO, and ZYU).

**Table S2. Summary of the culture media used in this study.**

**The 7 most effective culture media**

| Cuture medium (Abbreciation) | Source or Composition/L | Culture condition | Bacteria/Fungi |
|---|---|---|---|
| Brain Heart Infusion Agar + L-cysteine hydrochloride hydrate + hemin + vitamin K (BHI1) | BHI (CM1135B) +0.5 g/L L-cysteine hydrochloride hydrate (C121800) + 10 mg/L hemin (H811002) + 1 mg/L vitamin K (V884043) | Aerobic/ Anaerobic | Bacteria |
| Brain Heart Infusion Agar + colistin sulphate + naladixic acid + L-cysteine hydrochloride hydrate + hemin + vitamin K (BHI2) | BHI + 10 mg/L colistin sulphate (C9810-1) + 5 mg/L naladixic acid (N8878) + 0.5 g/L L-cysteine hydrochloride hydrate (C121800) + 10 mg/L hemin (H811002) + 1 mg/L vitamin K(V884043) | Aerobic/ Anaerobic | Bacteria |
| Actinomyces Isolation Agar (AIA) | 2.0g/L Sodium caseinate (M029018) + 0.1 g/L L-Asparagine (A105949) + 4.0 g/L Sodium propionate (V900330) + 0.5 g/L Dipotassium phosphate (P418694) + 0.1 g/L Magnesium sulfate (M2643) + 0.001 g/L Ferrous sulfate (F871869) + 20 g/L Agar (214010) | Aerobic/ Anaerobic | Bacteria |
| Chocolate Agar | From Becton Dickinson (BD) 228950 | Aerobic/ Anaerobic | Bacteria |
| Yeast extract-Casein hydrolysate Fatty Acids (YCFA) | From Solarbio LA4040 | Aerobic/ Anaerobic | Bacteria |
| Yeast Extract Peptone Dextrose (YPD) | 10 g/L Yeast Extract (212750), 20 g/L Peptone (211677), 20 g/L Glucose (G8270), 20 g/L Agar | Aerobic | Fungi |
| Czapek-Dox agar (CDA) | From PhytoTech Lab (C506) | Aerobic | Fungi |

**The other 23 culture media**

| Culture medium (Abbreciation) | Source or Composition/L | Culture condition | Bacteria/Fungi |
|---|---|---|---|
| Brain Heart Infusion Agar (BHI) | From Thermo Scientific CM1135B | Aerobic/ Anaerobic | Bacteria |
| Brain Heart Infusion Agar + colistin sulphate + naladixic acid (BHI-CN) | BHI + 10 mg/L colistin sulphate (C9810-1) + 5 mg/L naladixic acid (N8878) | Aerobic/ Anaerobic | Bacteria |
| Columbia Blood Agar with 5% Sheep Blood (CBA) | From Becton Dickinson (BD), Cat. No. 254071 | Aerobic/ Anaerobic | Bacteria |
| Tryptic Soy Broth (TSB) | BA-257107.06 | Aerobic/ Anaerobic | Bacteria |
| Bifidobacterium Selective Media (BSM) | Sigma 88517 | Aerobic/ Anaerobic | Bacteria |
| Cooked meat medium (BEEF) | 98 g/L Beef Heart (CB2514) + 20 g/L Proteose Peptone (P885893) + 2 g/L Dextrose (G8270) + 5 g/L Sodium Chloride (S5886) | Aerobic/ Anaerobic | Bacteria |

| | | | |
|---|---|---|---|
| Phenylethyl Alcohol Agar with 5% sheep's blood (PEA) | From Becton Dickinson (BD), 221739 | Aerobic/ Anaerobic | Bacteria |
| Colistin Naladixic Acid Agar with 5% sheep's blood (CNA) | From Becton Dickinson (BD), PA-254007.06 | Aerobic/ Anaerobic | Bacteria |
| Mannitol Slat Agar (MSA) | From Becton Dickinson (BD), 211407 | Aerobic/ Anaerobic | Bacteria |
| de Man Rogosa Sharpe Agar (MRS) | From Solarbio M8330 | Aerobic/ Anaerobic | Bacteria |
| Bacteroides Bile Esculin Agar (BBE) | From QDRS BIOTEC 12146 | Aerobic/ Anaerobic | Bacteria |
| MacConkey Agar (MAC) | From PD PM0801 | Aerobic/ Anaerobic | Bacteria |
| Deoxycholate Agar (DOC) | From Thermo Scientific CM0163B | Aerobic/ Anaerobic | Bacteria |
| D1 nutrient Agar (D1) | From Mast Group DM 179 | Aerobic/ Anaerobic | Bacteria |
| Kanamycin Vancomycin Laked Blood Agar (KVLB) | tryptic soy agar (22091) + 0.1 μg/mL kanamycin (A600286) + 7.5 μ g/mL vancomycin (S17059) + 10 μ g/mL vitamin K 1 + 0.05 ng/mL hemin + 5% laked horse blood (SR0048C) | Aerobic/ Anaerobic | Bacteria |
| Gifu anaerobic medium (GAM) | GMNB-GAM01 | Aerobic/ Anaerobic | Bacteria |
| Potato dextrose agar (PDA) | From Coolaber PM0520 + 10 g/L Yeast extract + 30 mg/L colistin + 30 mg/L vancomycin + 30 mg/L imipenem (MK0787) | Aerobic | Fungi |
| Sabouraud dextrose broth (SDB) | From Millipore 1.46366 + 5% Defibrinated sheep blood (P-62) + 5% rumen juice (A1101-01) + 20 g/L Agar | Aerobic | Fungi |
| Glycine-vancomycin polymyxin B agar | From BINDER AB113 | Aerobic | Fungi |
| Schaedler agar | Malt extract (218630), ox bile (B8631), oleic acid (O1008), glycerol (G8190), Tween 60 (P1629), colistin (30 mg/L), vancomycin (30 mg/L), and imipenem (30 mg/L) | Aerobic | Fungi |
| Malt agar | From Coolaber MM1051 | Aerobic | Fungi |
| Dixon agar | From HAIBO HB9217 + 30 mg/L Colistin, | Aerobic | Fungi |

| | | | |
|---|---|---|---|
| | 30 mg/L vancomycin, + 30 mg/L imipenem | | |
| CHROMagar Candida | From HY-WKM8001 | Aerobic | Fungi |

**Reference:**

Lagier J-C, Khelaifia S, Alou MT, Ndongo S, Dione N, Hugon P, et al. Culture of previously uncultured members of the human gut microbiota by culturomics. Nature Microbiology. 2016;1(12). doi: 10.1038/nmicrobiol.2016.203.

**Dataset S1. Information of the microbial species isolated from the mouse "sterile" organs by culturomics.**

**Dataset S2. The microbial species detected in the organs of at least three mice among 42 mice by culturomics.**

**Dataset S3. The dominate species isolated from different organs by culturomics.**

**Dataset S4. The microbial species detected by metagenomics assays.**

**Dataset S5. Taxonomic composition of microbes in each organ of the 10 mice detected by metagenomics assays.**
